# Comprehensive analysis of *Arabidopsis thaliana* DNA polymerase epsilon catalytic subunit A and B mutants – an insight into differentially expressed genes and protein-protein interactions

**DOI:** 10.1101/2022.02.14.480442

**Authors:** Anushka M. Wickramasuriya, Thulani M. Hewavithana, Kithmee K. de Silva, Ihsan Ullah, Jim M. Dunwell

**Affiliations:** Department of Plant Sciences, Faculty of Science, University of Colombo, Colombo 03, Sri Lanka; School of Agriculture, Policy and Development, University of Reading, UK

## Abstract

One of the main replicative enzymes in most eukaryotes, DNA polymerase ε (POLE), is composed of four subunits, namely a single catalytic and three regulatory subunits. In Arabidopsis, the catalytic subunit of POLE is encoded by two genes: *Arabidopsis thaliana DNA POLYMERASE EPSILON CATALYTIC SUBUNIT A* (*AtPOL2A*) and *B* (*AtPOL2B*). Although studies have shown *AtPOL2A* to be involved in various biological processes, the role of *AtPOL2B* is unclear. Here, we investigated the transcriptomes of both *atpol2a* and *atpol2b* mutants, and the promoter sequences to provide a better insight into the targets of *AtPOL2s* at the molecular level. In the present study, leaf cDNA libraries of four *AtPOL2* mutants (*atpol2a-1* and *atpol2b-1, -2* and *- 3*) were sequenced using the Illumina platform. Analysis of gene expression profiles identified a total of 198, 76, 141 and 67 differentially expressed genes in *atpol2a-1*, *atpol2b-1*, *atpol2b-2* and *atpol2b-3*, respectively; the majority of pericentromeric transposable elements were transcriptionally active in *atpol2a-1* as compared to *atpol2b* mutants and wild type. Protein-protein interaction network analysis and molecular docking identified three (CER1, RPA1E and AT5G60250) and two (PR1 and AT5G48490) proteins as potential interactors (*cluster size > 60* and *balanced score < -900*) of AtPOL2A and AtPOL2B, respectively; Interestingly, these five proteins also showed a significant interaction between POLE catalytic subunit of *Saccharomyces cerevisiae.* Our *in silico* promoter analysis showed that the *AtPOL2A* promoter sequence is overrepresented with *cis*-acting regulatory elements (CREs) associate with cell cycle regulation, meristematic/reproductive tissue-specific pattern of expression and MYB protein recognition, whereas the *AtPOL2B* promoter sequence was mainly enriched with stress-responsive elements. The information provided here has led to the identification of targets of AtPOL2s at the molecular level and CREs putatively associated with the regulation of *AtPOL2*s. To our knowledge, this study provides the first comparative transcriptome profiling of single-gene mutants of *AtPOL2s*.

## Introduction

In all living organisms, DNA polymerases (DNA Pols) play a central role in the regulation of DNA replication and repair to ensure faithful transmission of genetic information. Eukaryotic DNA Pols have been arranged into five well characterized families namely A, B, C, D and X [1–3]; of these, B, C and D family members are the main regulators of nuclear DNA replication [2]. In most eukaryotes, the members of the family B DNA Pols ‒ α, δ and ε are the main replicative DNA Pols; knock out mutants of the main catalytic subunit of DNA polymerase ε (POLE), for example, *tilted* (*til*) *1-1* to -*3* in Arabidopsis are embryo lethal [4, 5]. The present study focuses on the catalytic subunit of the Arabidopsis POLE complex that has been shown to have diverse regulatory roles, for example in DNA replication and repair, and epigenetic gene silencing.

The POLE catalytic subunit is known to be involved in both replisome assembly and leading strand synthesis in budding yeast [6–8]. It is characterized by the presence of a catalytically active N-terminal and an inactive C-terminal domain. The active sites for polymerase and exonuclease activities are found within the N-terminal domain, the domain which is highly conserved among other family B polymerases i.e. DNA polymerase α, DNA polymerase β and bacteriophage DNA polymerases (T4 and RB69) [9, 10]. Recently it has been found that POLE maintains the functional integrity of the replication fork through phosphorylation of the serine 430 residue on the catalytic subunit [11].

Although a larger portion of the POLE catalytic subunit accounts for the C-terminal domain [10], with the exception of the DIE motif, this domain lacks catalytic residues for both polymerase and exonuclease activities [12]. Tahirov and co-workers have reported that the presence of catalytically active and inactive domains in POLE is due to a more complex evolutionary scenario than a simple tandem duplication [12]. Based on their findings, one possible hypothesis for the existence of this large catalytically inactive domain would be the presence of at least two family B DNA polymerases and a DNA polymerase δ in the archaeal ancestor of the family B DNA polymerases. However, during evolution, selection pressure may have favored the existence of the catalytically inactive large C-terminal domain of POLE as a means of ensuring the structural integrity of the POLE complex [12].

In *Arabidopsis thaliana*, the catalytic subunit of POLE is encoded by two genes [4, 13], *AtPOL2A* (also known as *POLE1*, *TIL1*, *ABSCISIC ACID OVERSENSITIVE* 4 (*ABO4*), or *EARLY IN SHORT DAYS 7* (*ESD7*); TAIR locus id: AT1G08260) and *AtPOL2B* (also called *POLE1B*; TAIR locus id: AT2G27120). The *AtPOL2A* gene is approximately 16 kb long (49 exons) and encodes a protein of 2161 amino acids, whereas the approximately 13 kb long *AtPOL2B* (49 exons) encodes a protein of 2138 amino acids which is 78.5% identical to AtPOL2a [4,13–15]. Both AtPOL2A and B contain all the motifs that are necessary for a functional POLE catalytic subunit.

Gene expression studies have detected *AtPOL2A* transcripts in a range of plant tissues i.e. actively growing tissues (floral meristem and flowers until anthesis) and mature tissues (5-week old leaves) but at a relatively low level [4]. However, the expression of *AtPOL2B* has been detected mostly under adverse environmental conditions and loss of function of *AtPOL2B* has resulted in no visible phenotype, suggesting that *AtPOL2A* is the main catalytic subunit encoding gene [13]. Over the past, four hypomorphic alleles of *AtPOL2A* (designated as *til1-4*, *abo 4-1* and *abo4-2*, and *esd7-1*) have been identified and characterized. The *til1-4* mutant lines, which carry two G to A mutations in exon 12 and intron 14 have exhibited longer cell cycles during embryo development, delayed development and larger cells [4]; both *abo4-1*, which has a point mutation that substitutes Glycine to Arginine (at the 534 residue) and *abo4-2,* which harbors a T-DNA insert in the 12^th^ exon, have shown sensitivity to abscisic acid (ABA), enhanced homologous recombination, sensitivity to DNA damage agents and constitutive expression of DNA repair genes [14]; the *esd7-1*, which has a Glycine to Arginine in amino acid position 992, has displayed early flowering and altered growth [15]. Although the molecular function of *AtPOL2B* is not well documented, detection of developmental defects in embryos and early flowering phenotypes in double mutants of *AtPOL2*genes have led to the discovery of partial functional redundancy of these genes during embryogenesis [4] and vegetative to floral transition [15].

Transcriptome analysis on *abo4-1* has shown that several genes related to DNA replication and repair, cell cycle control and flower development are differentially expressed in the mutant as compared to that of wild-type (WT) plants. Although *AtPOL2A* has been functionally characterized, functional characterization of its isoform is hampered due to the functional redundancy of the two genes. Hence, in the present study, we explore the effect of disrupting *AtPOL2A* and *AtPOL2B* genes on whole-transcription to gain insight into potential downstream target genes of both *AtPOL2A* and *B* genes and their interactions at a molecular level.

## Results

### Molecular confirmation of *atpol2* mutants

The homozygous/heterozygous nature of T-DNA mutant lines and the orientation of T-DNA insert(s) were confirmed through a PCR-based screening method. We found that the homozygous, viable *atpol2a-1* line harbours at least two T-DNA inserts in opposite orientations within the *AtPOL2A* locus (S1 Fig). It was also found that one T-DNA insert is positioned within chromosome 1, after the 2,594,915^th^ base (5’—[left border (LB) — right border (RB)]—3’ orientation) and the second insert is positioned just before the 2,594,924^th^ base in the 5’—[RB — LB]—3’ orientation. Although the Salk Institute Genomic Analysis Laboratory (SIGnAL) T-DNA express: *Arabidopsis* gene mapping tool (http://signal.salk.edu) supports the presence of an insert at the 12^th^ intron in 5’—[LB — RB]—3’ orientation in the *atpol2a-1* mutant, the additional insert found within intron 12 of the *AtPOL2A* has not been reported previously.

Similarly, the individuals of each mutant line of *AtPOL2B* used in the present study were (*atpool2b*-*1* to -*3*) confirmed by PCR amplification (S2 Fig). Consistent with the mutant information provided in the *Arabidopsis* genome browser and SIGnAL T-DNA express: *Arabidopsis* gene mapping tools, we found that *atpol2b-1* homozygous mutant line harbours a T-DNA insert in the second intron (307 bp from the transcription start site (TSS)) in the 5’—[RB — LB]—3’ orientation. Furthermore, in the *atpol2b-3* line, the insertion was in exon 34 (8930 bp from the TSS). However, the *atpol2b-2* line exhibited the presence of at least two T-DNA inserts within *AtPOL2B*; one insert was oriented in the 5’—[LB — RB]—3’ direction (4,408 bp from the TSS) and the second insert was detected after 4,429 bp from the TSS in a 5’—[RB — LB]—3’ orientation within intron 15.

### Morphology of *atpol2* mutant lines

The homozygous *atpol2a-1* line showed an apparent mutant phenotype as compared to WT plants. These mutants were early flowering when grown under a 16 h photoperiod (Fig 1A). Additionally, these individuals produced considerably smaller siliques (0.85 ± 0.02 cm; Fig 1B) with a reduced number of seeds (16 ± 1.63 seeds/ silique) as compared to WT plants (58 ± 1.45 seeds/ silique; (Fig 1C)). However, none of the *atpol2b* mutants examined here showed any visible phenotypic variation as compared to WT plants, under our experimental conditions (Fig 1A-B).

**Fig 1.**
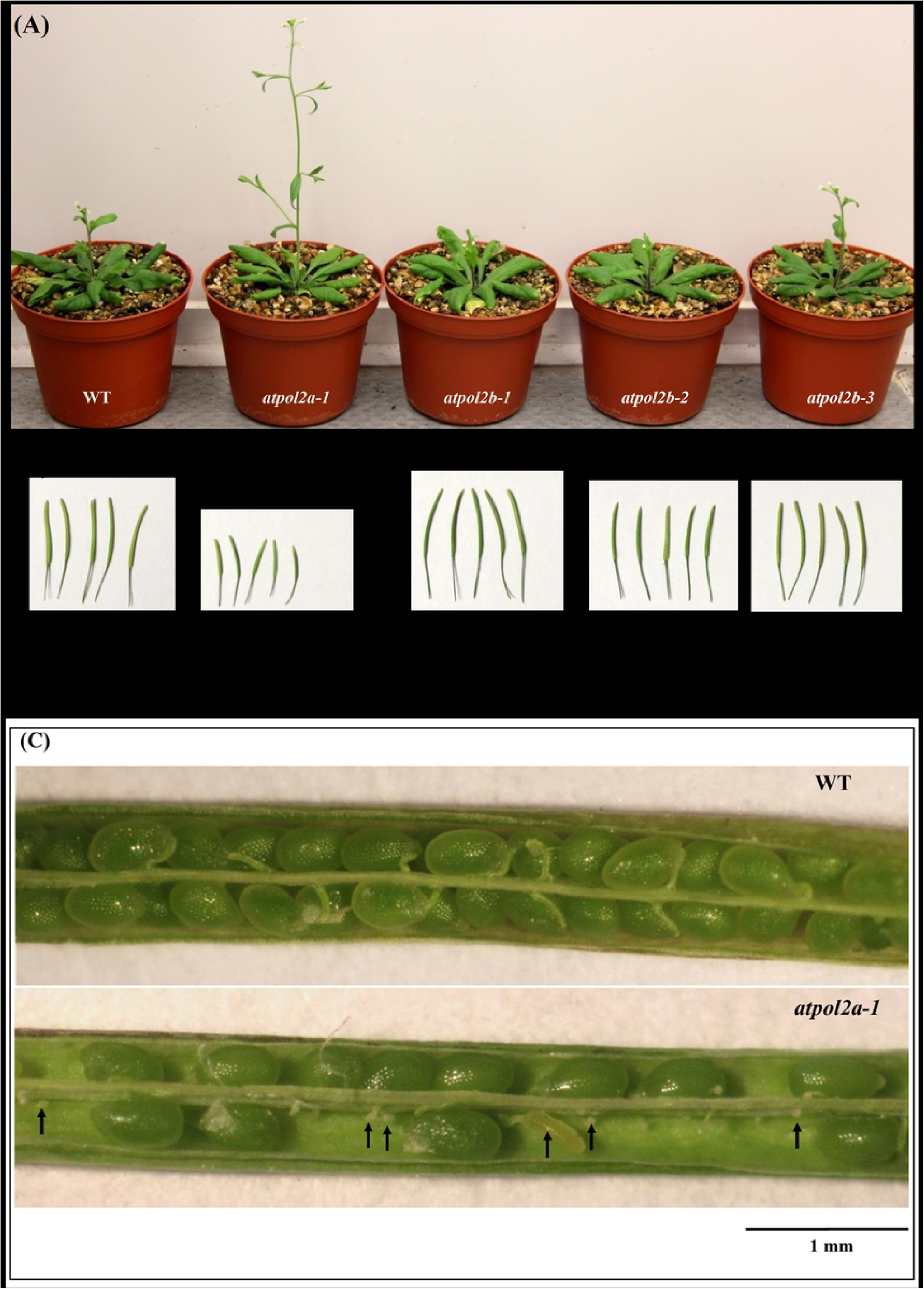
Morphological characters of WT and homozygous *atpol2* mutant plants when grown under a 16h photoperiod. (A) 26 d old plants. (B) mature siliques. (C) opened siliques of a WT plant having normal green seeds and a homozygous *atpol2a-1* mutant with aborted seeds (black arrows). The opened siliques of all three *atpol2b* lines were similar to those of WT plants and therefore are not shown here.

### Expression of *AtPOL2A* and *B* genes in *atpol2a-1* and *atpol2b-1* to *-3* mutant lines

In the present study, we used the Illumina platform to sequence cDNA libraries constructed from leaf tissues of WT and homozygous insertional lines of *AtPOL2* genes. A detailed analysis of transcriptome data derived from this study is given in the latter sections. Only the expression levels of the *AtPOL2A* and *B* genes in their respective mutant lines are considered here.

Visualization of the reads mapped to the *AtPOL2A* locus in the *atpol2a-1* derived transcriptome clearly showed increased transcription from the regions up and downstream of the T-DNA insertion site as compared to the WT. Furthermore, analysis of the reads mapped to the *AtPOL2B* locus in transcriptomes derived from *atpol2b-1* to *-3* mutant lines showed that more reads are mapped to regions downstream of the insertion; there were none mapped upstream of the insertion site.

To further confirm the expression patterns detected for *AtPOL2A* and *B* in their mutant lines, RT-qPCR analysis was performed using two sets of primers for each gene: one primer pair was designed from a region upstream to the insertion site and another pair from a region downstream of the insertion. This further confirmed that the *atpol2a-1* mutant line produces wild-type *AtPOL2A* transcripts from regions both up and downstream of the T-DNA insertion site (Fig 2A). Additionally, all the *AtPOL2B* mutant lines studied here showed notable expression of *AtPOL2B* transcripts downstream of the T-DNA insertion as compared to the WT (Fig 2B).

**Fig 2.**
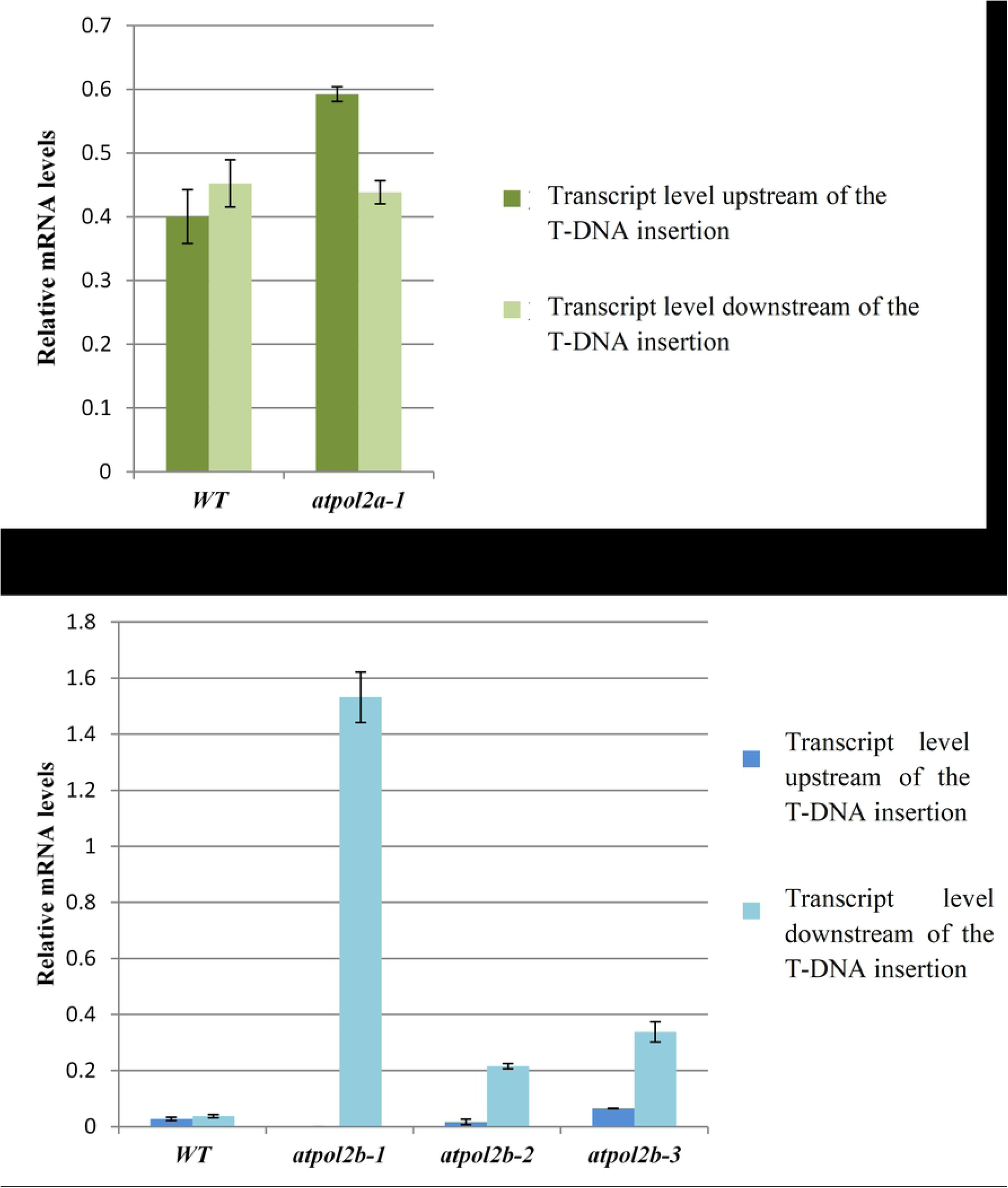
Expression levels of *AtPOL2A* and *B* transcripts in WT and their mutant lines detected through RT-qPCR. (A) Expression of *AtPOL2A* in leaf tissues of WT and *atpol2a-1*. (B) Expression of *AtPOL2B* in leaf tissues of WT, *atpol2b-1* to *-3*.

### Transcriptome analysis of homozygous *atpol2* mutants

Genome-wide transcriptome analysis using RNA-sequencing (RNA-Seq) was performed on homozygous *atpol2a-1* and *atpol2b-1* to *-3* mutant lines to identify potential downstream targets of *AtPOL2* genes. In total, 179025732, 167015310, 180266730, 158718412 and 223844382 reads were obtained from the leaf cDNA libraries of the WT, *atpol2a-1*, *atpol2b-1*, *atpol2b-2* and *atpol2b-3*, respectively. Approximately, 98% of sequence reads were aligned to the *A. thalaiana* reference genome (TAIR 10).

In total, the present study detected transcript abundance of 27,409 annotated transcripts in at least one of the sequencing libraries examined; the majority of transcripts were protein-coding. In addition, a considerable fraction of transposable elements (TEs) was transcriptionally active in the *atpol2a-1* mutant (Fig 3).

**Fig 3.**
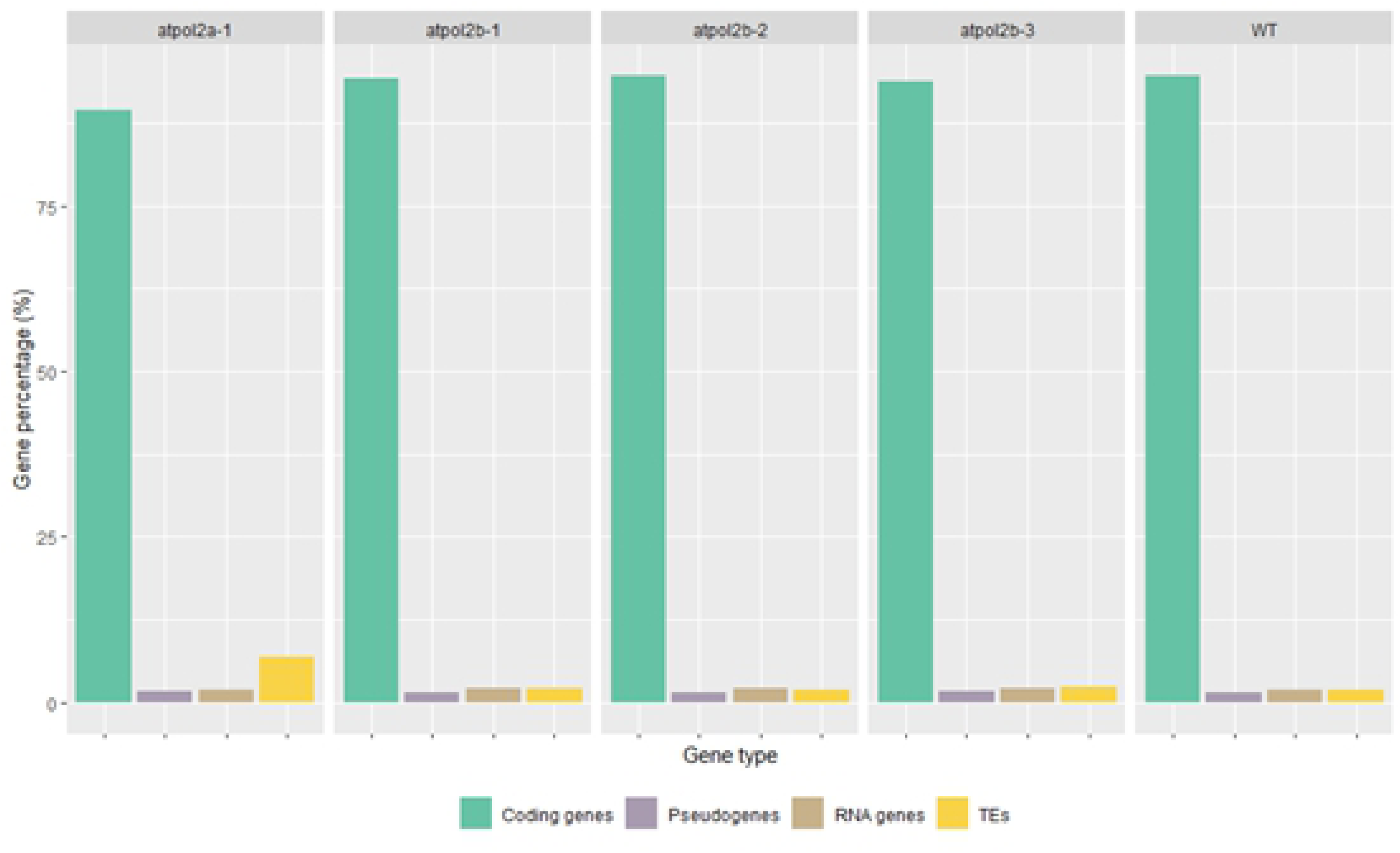
Different gene types detected in each sequencing library. Only the results obtained through the Cufflinks package are shown here.

Differential gene expression analysis was performed using three computational tools based on the negative binomial distribution to generate a high-confidence list of differentially expressed genes (DEGs). As expected there was a variation in the DEGs detected through the three computational methods (Fig 4); DEGs commonly identified through all three analysis tools (Cuffdiff, DESeq and edgeR) for each mutant line were considered for the downstream functional analysis. It was evident that at least 198 genes in *atpol2a-1*, 76 genes in *atpol2b-1*, 141 genes in *atpol2b-2* and 67 genes in *atpol2b-3* are differentially expressed as compared to WT (S1 Table). Further comparison of the DEGs detected in *atpol2a-1* in the present study and DEGs reported previously for *AtPOL2A* mutants [16, 17] showed that at least 11 genes are commonly differentially expressed in all *AtPOL2A* gene mutants (Fig 5; Table 1).

**Fig 4.**
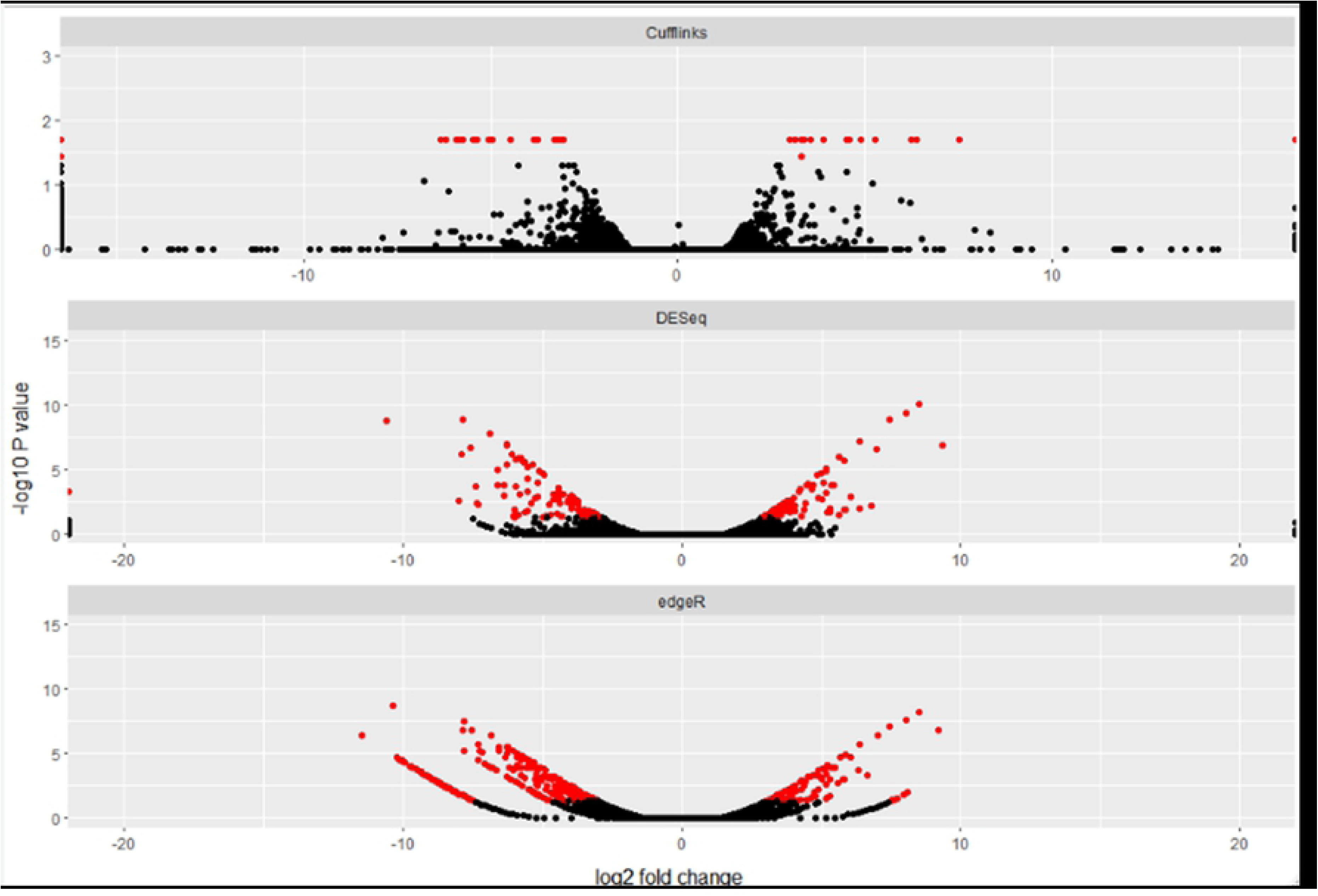
Volcano plots showing the differential gene expression. DEGs detected in *atpol2a-1* mutants as compared to WT are shown here. Genes with an adjusted *p-value* of ≤ 0.05 and -1 ≥ *log_2_* fold-change ≥ 1 are shown as red dots.

**Fig 5.**
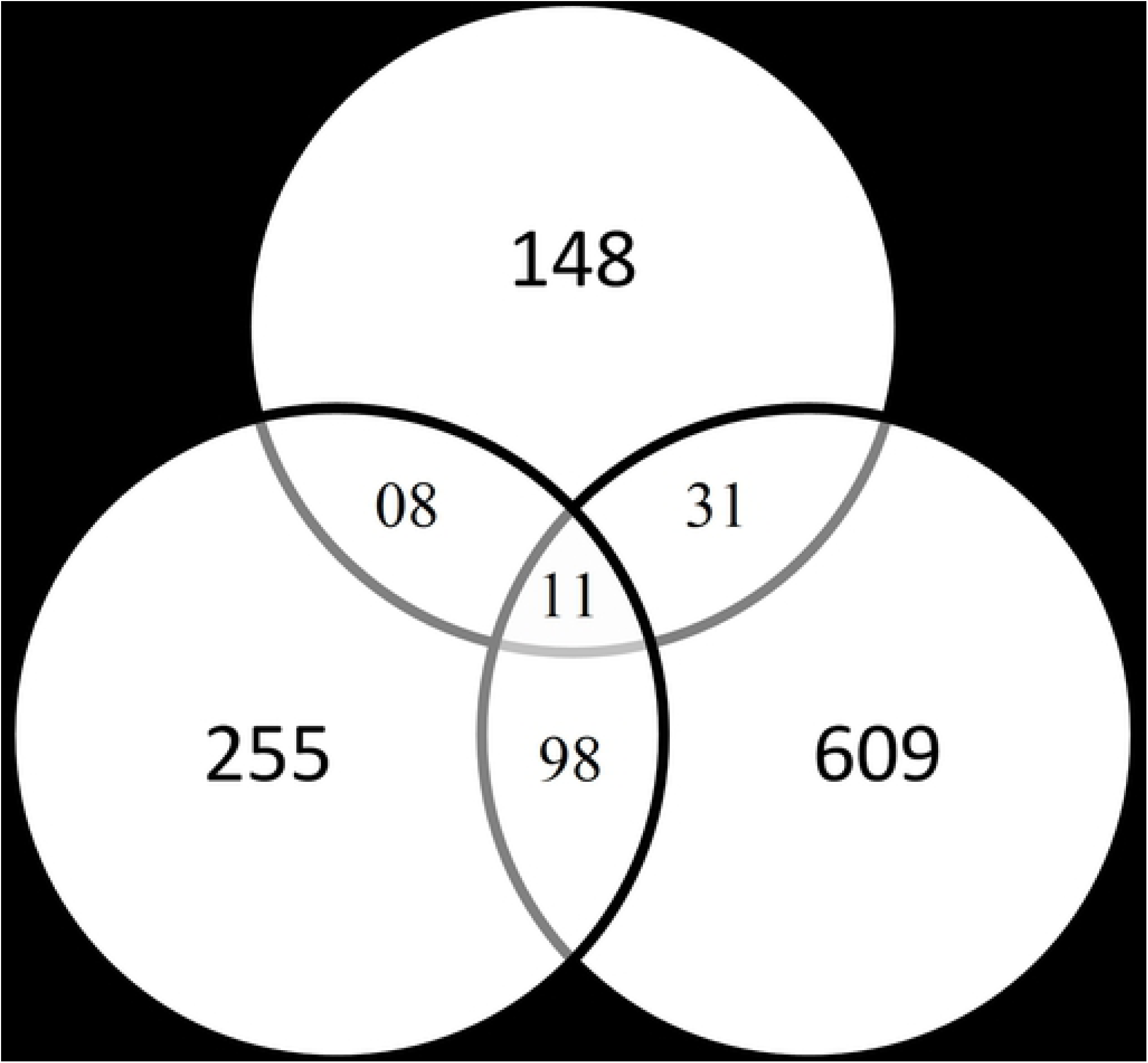
Venn diagram depicting the DEGs detected in different *AtPOL2A* mutants.

**Table 1.**
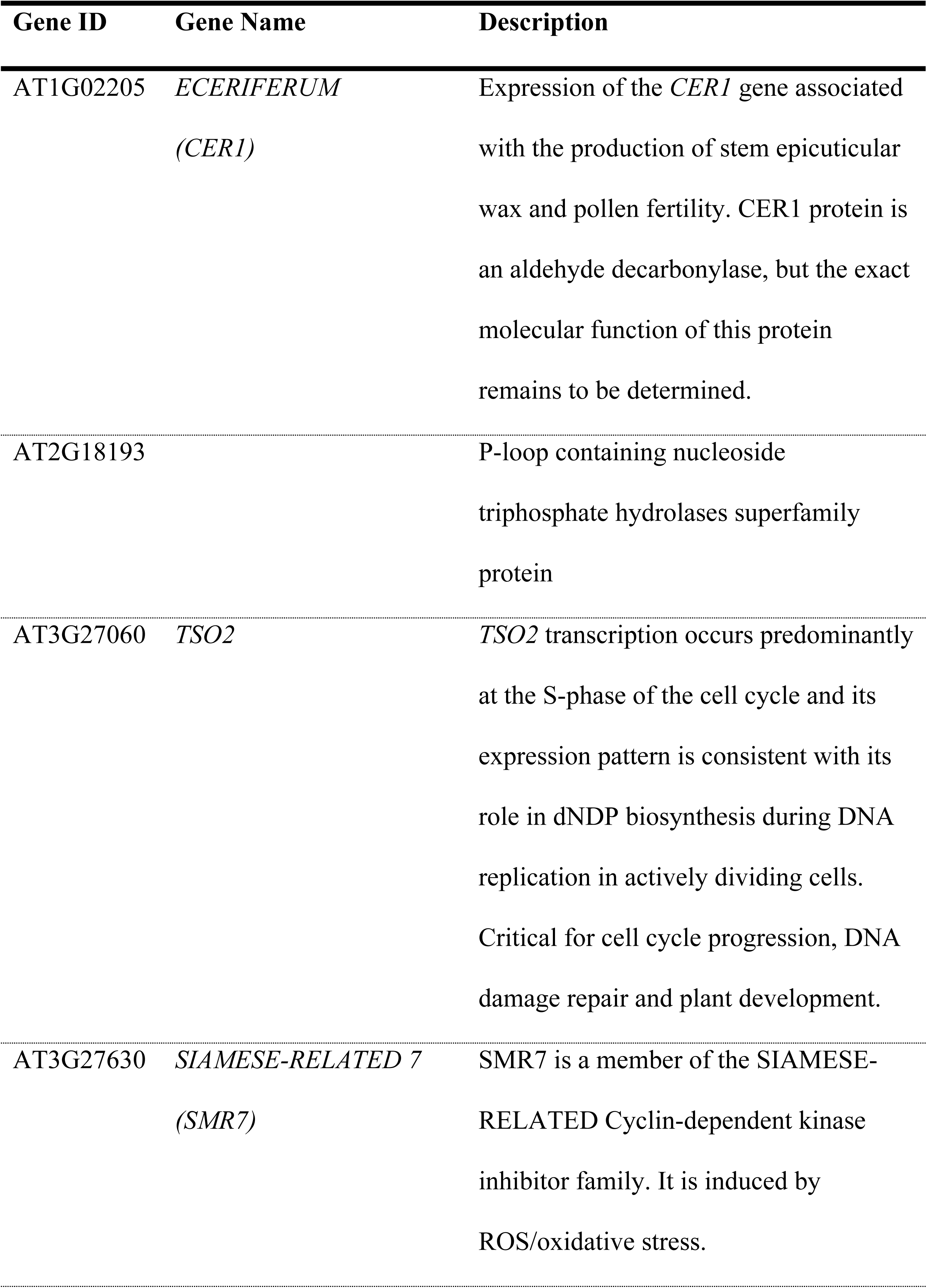
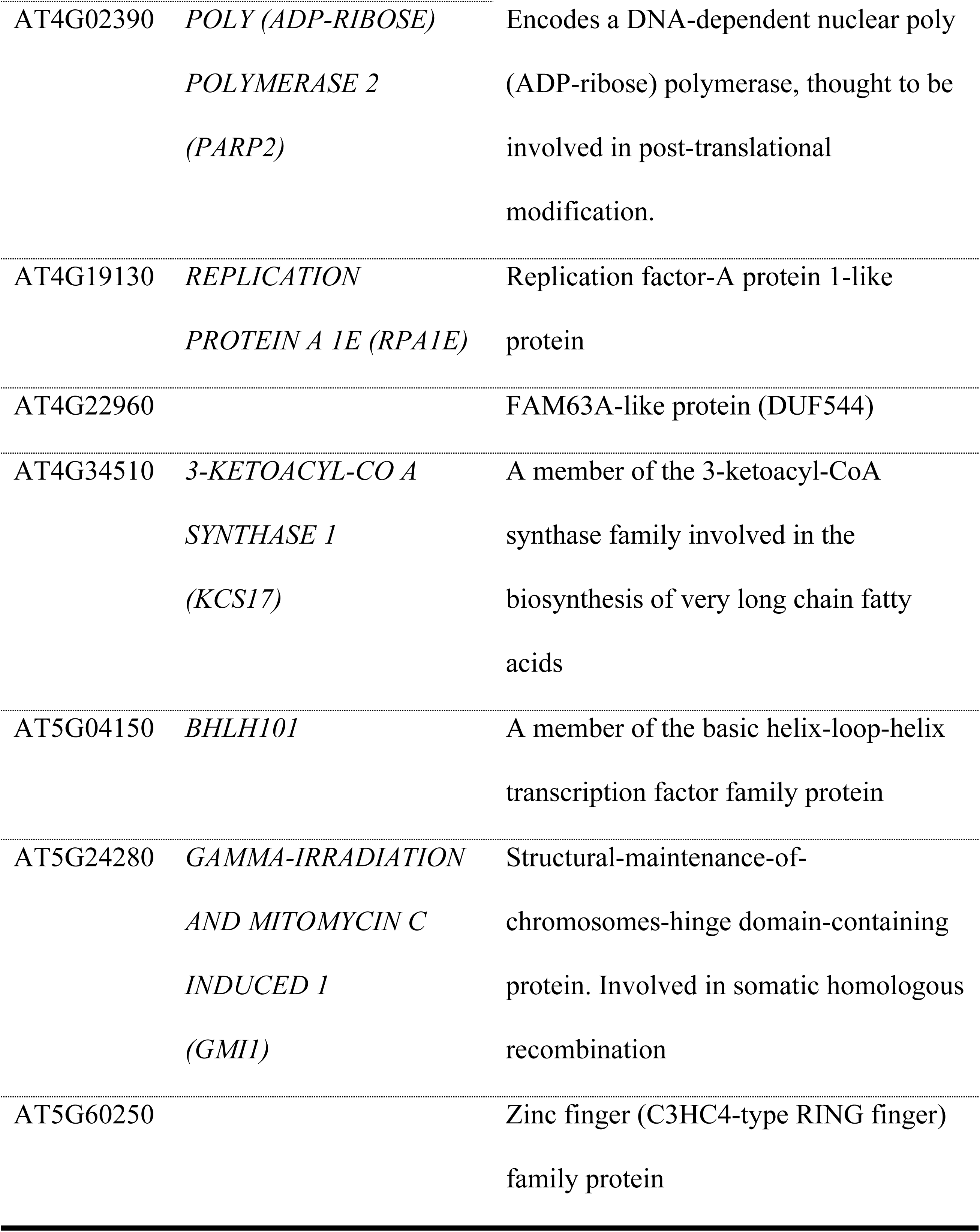
List of genes differentially expressed in *AtPOL2A* gene mutants (*atpol2a-1* and *abo4*).

Gene Ontology (GO) enrichment analysis of the 198 DEGs detected in *atpol2a-1* showed significant enrichment (P-value ≤ 0.05) for GO terms such as response to defense, oxidation-reduction process, response to UV-A, response to light stimulus, response to water deprivation, response to ABA and response to jasmonic acid; the list of enriched GO terms is shown in Fig 6. Analysis of the promoter sequences (1000 bp upstream genomic region from the TSS) of all DEGs detected in *atpol2a-1* using the Multiple Em for Motif Elicitation (MEME) suite showed five motifs are significantly overrepresented in DEGs (P < 0.05) (Table 2). The possible biological roles of these motifs were studied using Gene Ontology for Motifs (GOMo) tool. It was noted that several motifs possibly function in regulating circadian rhythm, and responses to ABA and jasmonic acid. Interestingly, a motif (motif-3: WCMYBYYCTCTYCTYTCYYY) that may function in regulating megagametogenesis was discovered in promoter regions of 28 DEGs. Further analysis of these discovered motifs using PlantPAN 3.0 showed that motif-1 and 2 may interact with MADS-box transcription factor SUPPRESSOR OF OVEREXPRESSION OF CO1 (SOC1)/AGAMOUS-LIKE 20 (AGL 20) [AT2G45660], whereas motif-4 may interact with ABA INSENSITIVE 5 (ABI5; AT2G36270) and/or PHYTOCHROME INTERACTING FACTOR 1 (PIF 1; AT2G20180).

**Fig 6.**
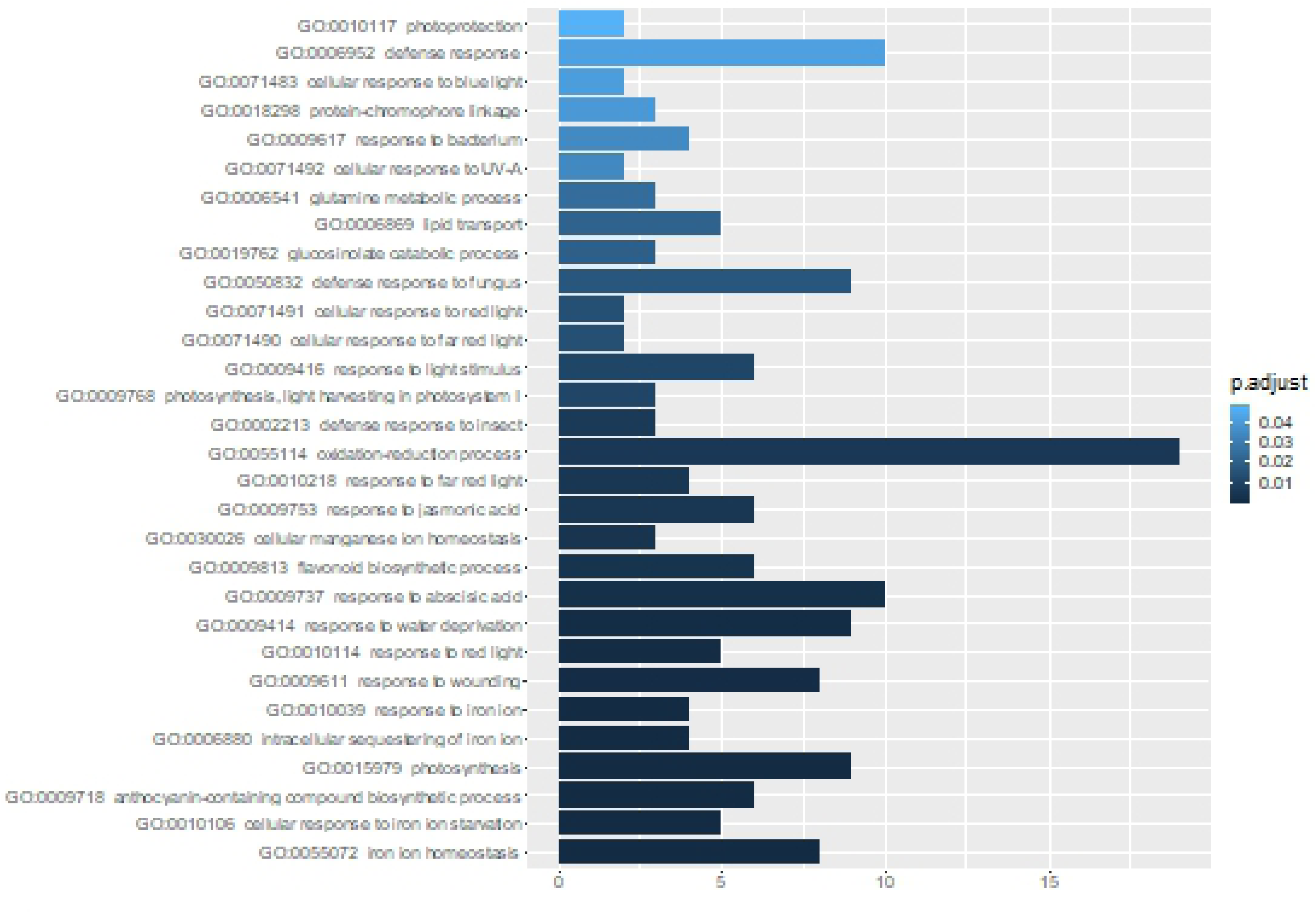
GO enrichment analysis of DEGs detected in *atpol2a-1*. Only the GO terms related to biological processes at P-value ≤ 0.05 are shown here.

**Table 2.**
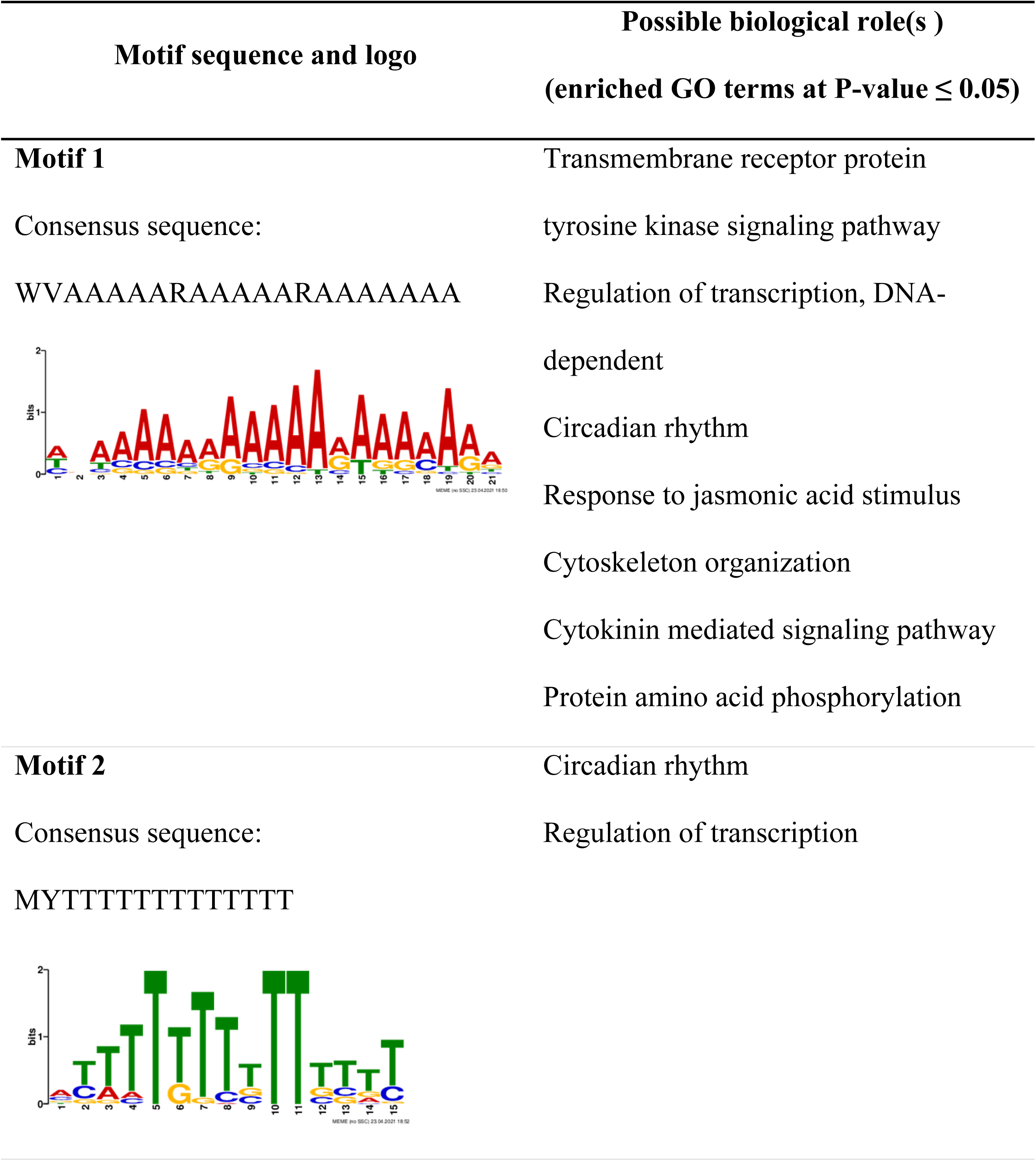
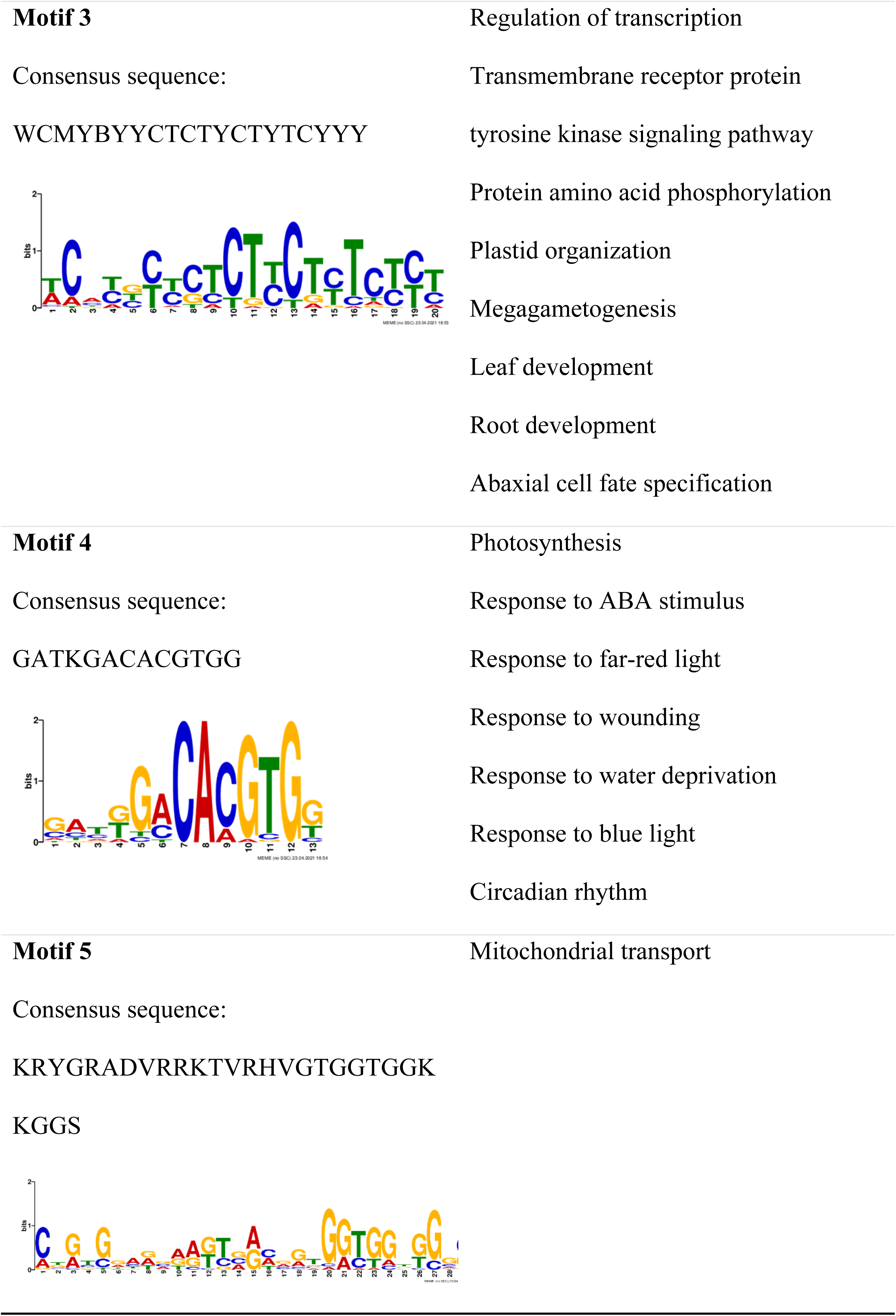
Motifs discovered in promoter regions of DEGs detected in *atpol2a-1*.

| Motif sequence and logo | Possible biological role(s )<br>(enriched GO terms at P-value ≤ 0.05) |
| --- | --- |
| **Motif 1**<br>Consensus sequence:<br>WVAAAAARAAAAARAAAAAA | Transmembrane receptor protein<br>tyrosine kinase signaling pathway<br>Regulation of transcription, DNA-dependent<br>Circadian rhythm<br>Response to jasmonic acid stimulus<br>Cytoskeleton organization<br>Cytokinin mediated signaling pathway<br>Protein amino acid phosphorylation |
| **Motif 2**<br>Consensus sequence:<br>MYTTTTTTTTTTTTTT | Circadian rhythm<br>Regulation of transcription |
| **Motif 3** | Regulation of transcription |
| Consensus sequence: | Transmembrane receptor protein |
| WCMYBYYCTCTYCTYTCYYY | tyrosine kinase signaling pathway |
|  | Protein amino acid phosphorylation |
|  | Plastid organization |
|  | Megagametogenesis |
|  | Leaf development |
|  | Root development |
|  | Abaxial cell fate specification |
| **Motif 4** | Photosynthesis |
| Consensus sequence: | Response to ABA stimulus |
| GATKGACACGTGG | Response to far-red light |
|  | Response to wounding |
|  | Response to water deprivation |
|  | Response to blue light |
|  | Circadian rhythm |
| **Motif 5** | Mitochondrial transport |
| Consensus sequence: |  |
| KRYGRADVRRKTVRHVGTGGTGGK |  |
| KGGS |  |

Comparison of DEGs detected in the three *atpol2b* mutants (*atpol2b*-*1* to -*3*) showed a substantial difference among the mutants (Fig 7); only 12 genes were commonly differentially expressed in all three mutants and their descriptions are given in Table 3. Genes that were differentially expressed in at least two of the *atpol2b* studied here were considered as DEGs in *AtPOL2B* mutants and used in downstream analyses.

**Fig 7.**
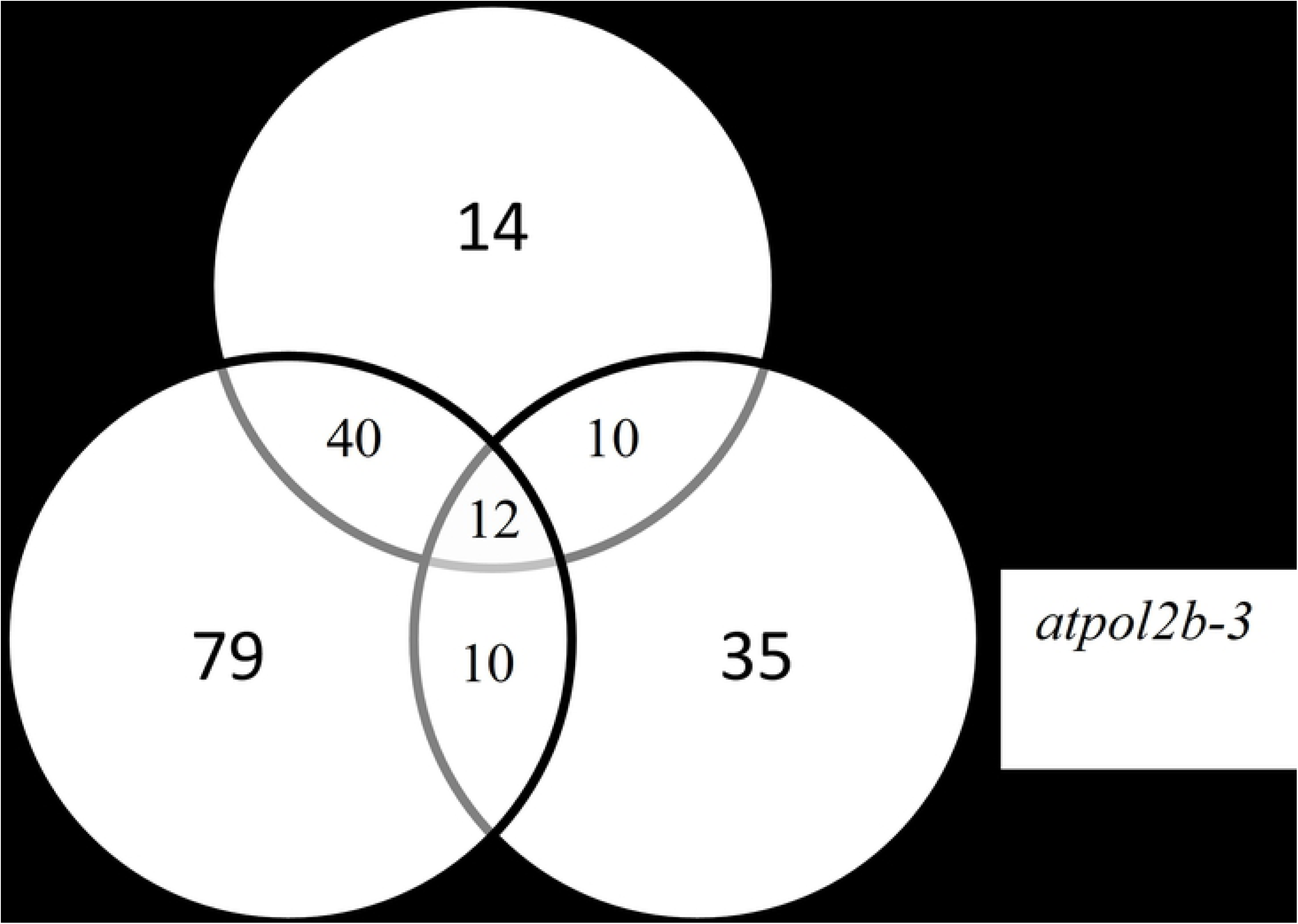
Venn diagram depicting the DEGs detected in *atpol2b1* – *3*.

**Table 3.**
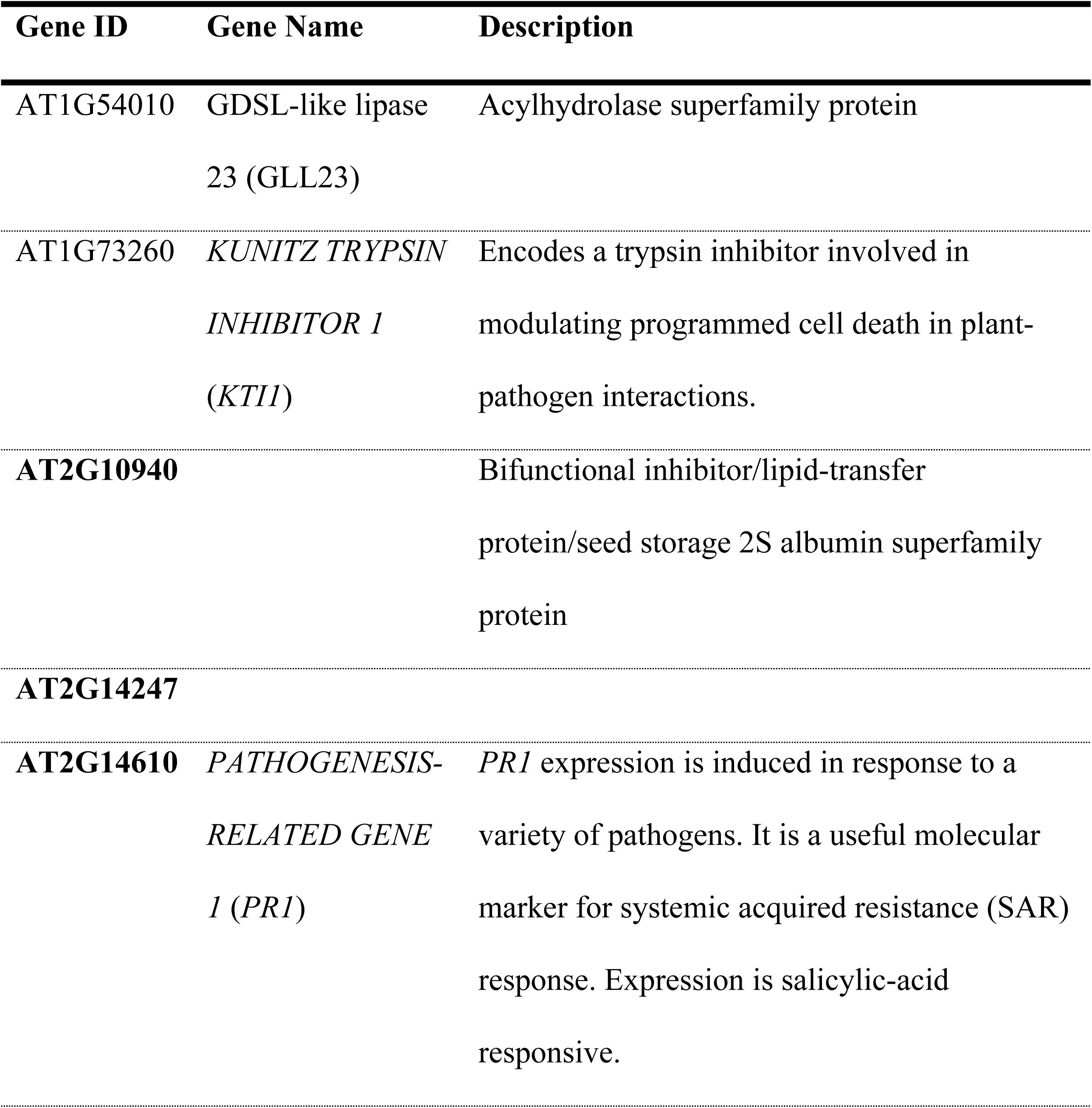
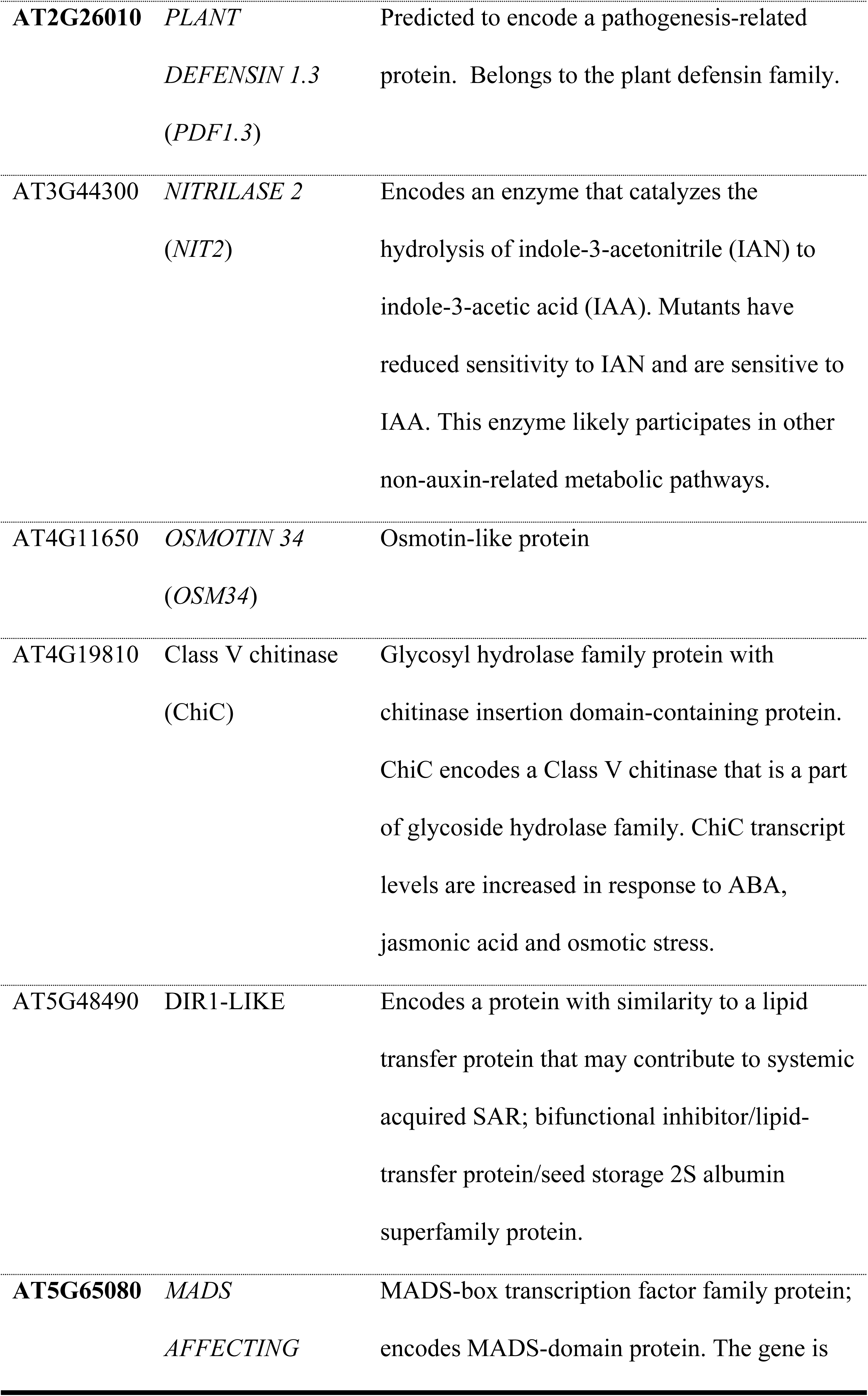
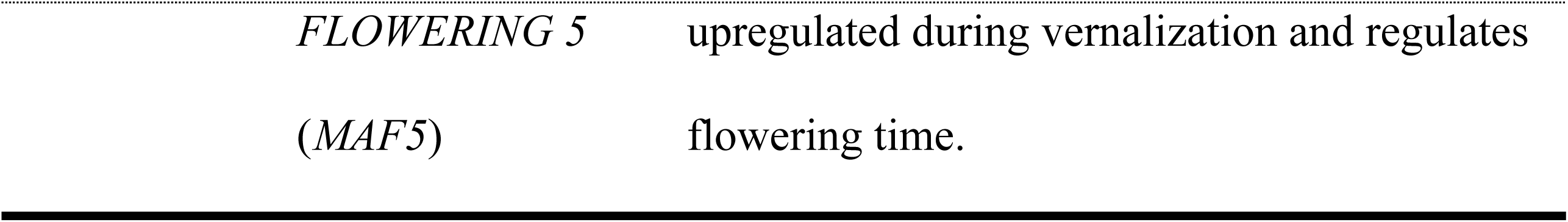
List of DEGs commonly detected in *atpol2b* mutants. Bold gene ids designate genes that showed differential expression in both *atpol2a* and *b* mutants.

GO enrichment analysis of DEGs detected in *atpol2b* mutants revealed a significant enrichment (P-value ≤ 0.05) for GO terms relating to response to defense, induced systemic resistance, response to salt stress, lipid transport and iron homeostasis (Fig 8).

**Fig 8.**
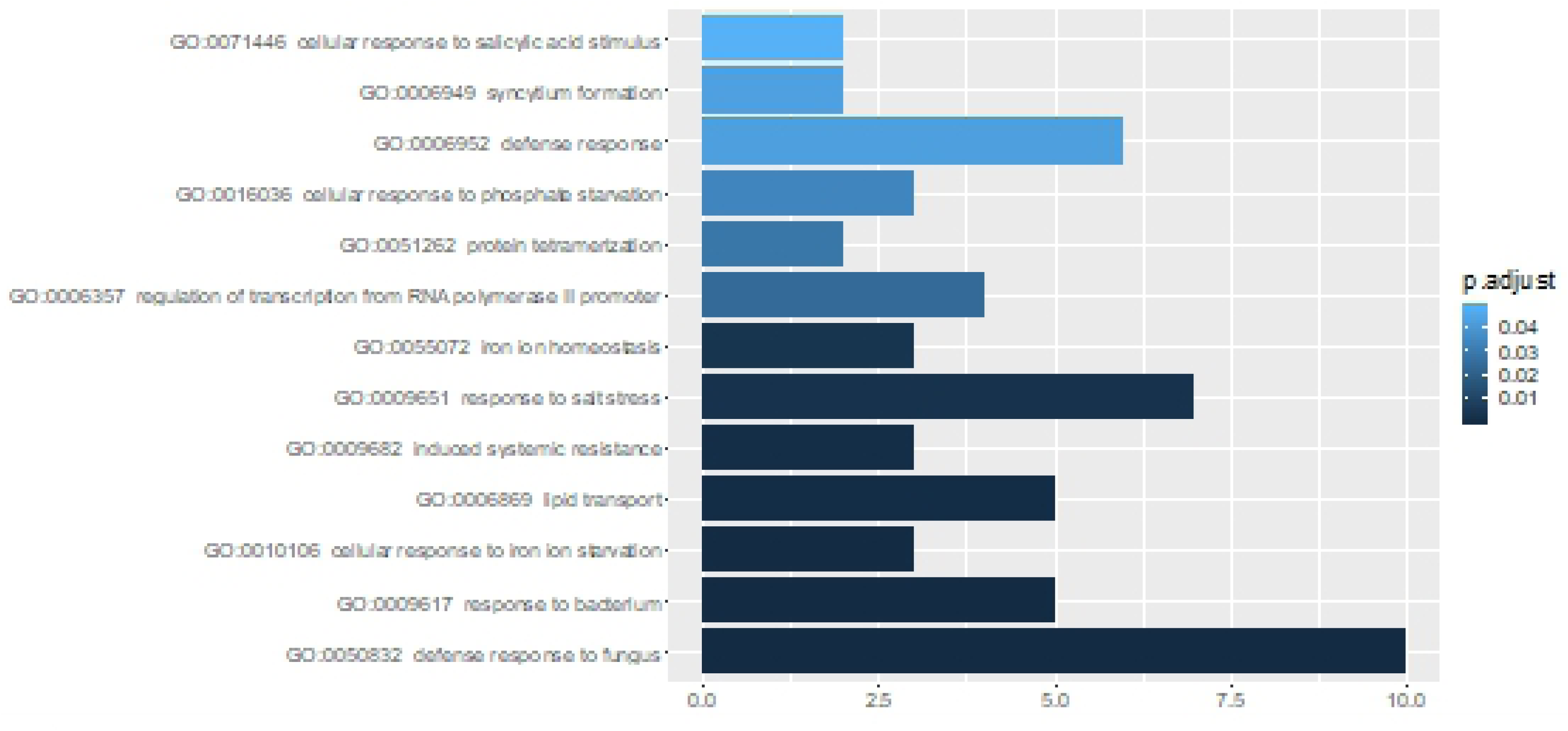
GO enrichment analysis of DEGs detected in at least two *atpol2b* mutants. Only the GO terms related to biological processes at P-value ≤ 0.05 are shown here.

Analysis of the promoter sequences (1000 bp upstream from the TSS) of DEGs identified in at least two *atpol2b* mutants revealed enrichment of five motifs (Table 4). The majority of DEGs contained motifs I (68/72 genes) and IV (66/72 genes) in their promoter regions that may interact with MADS-box transcription factor SOC1/ AGL 20.

**Table 4.**
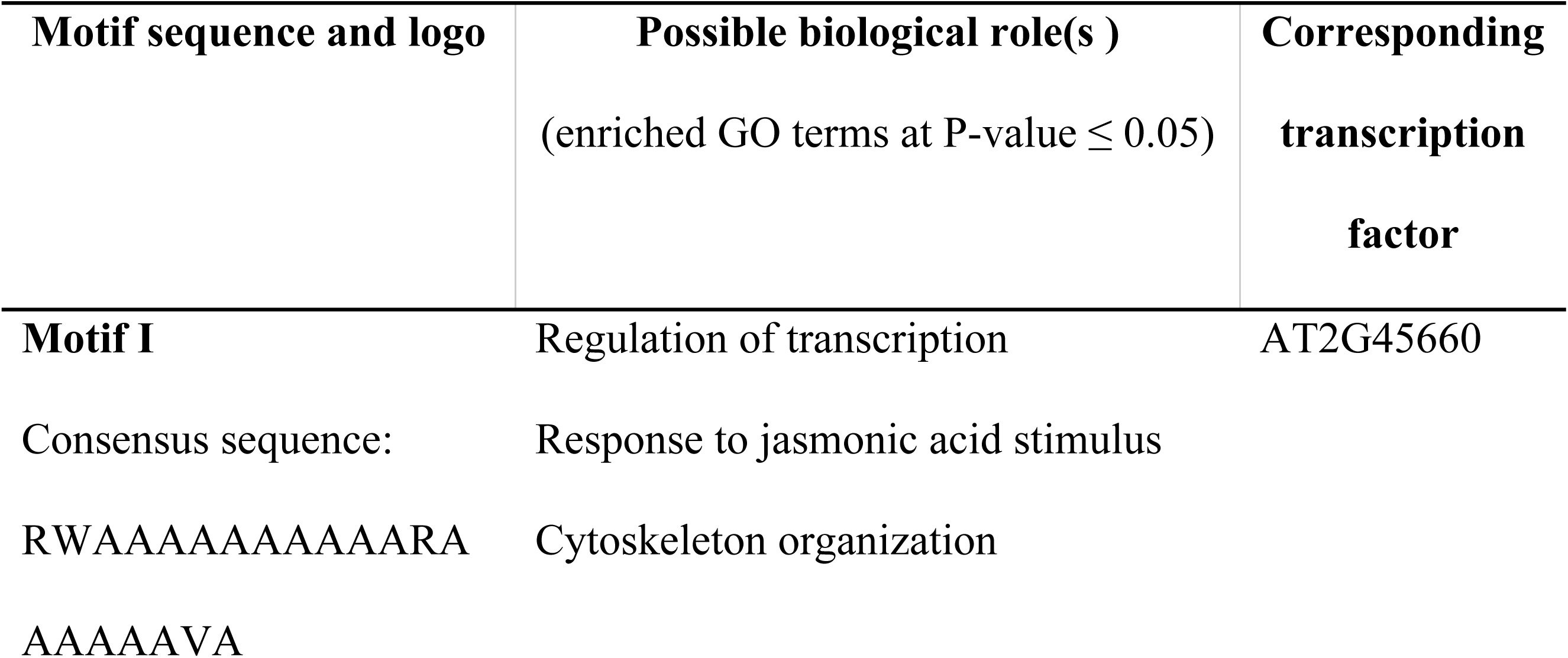
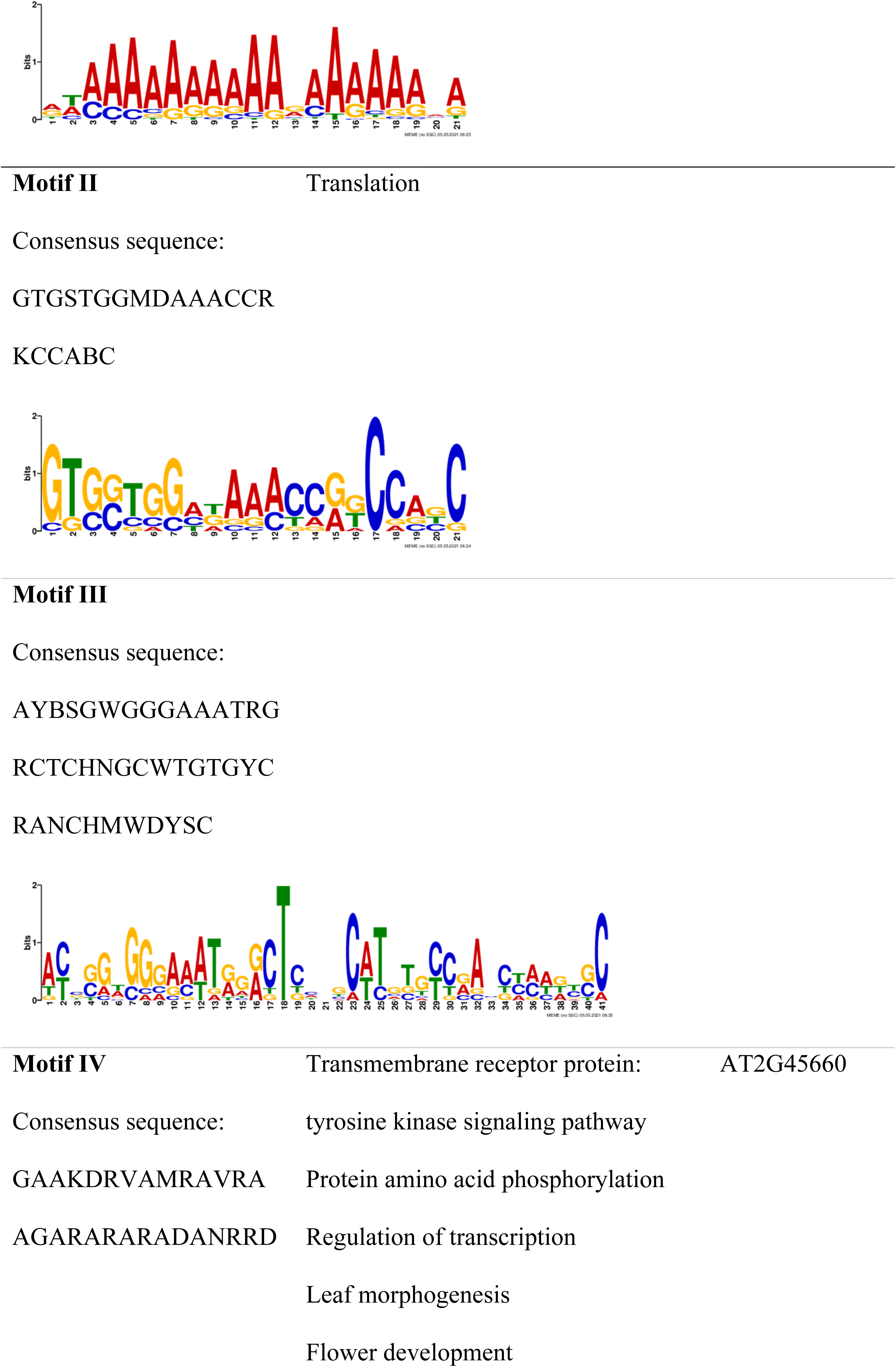
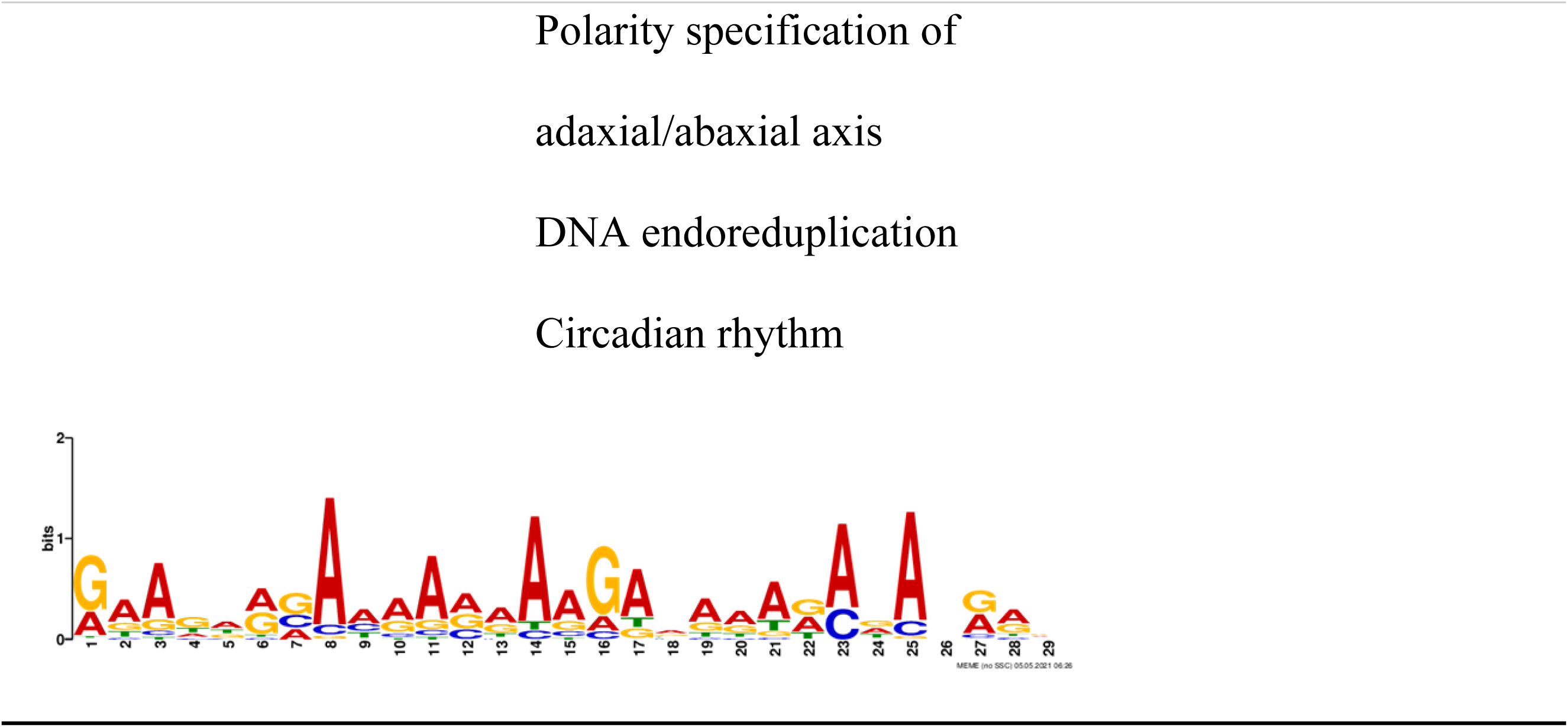
Motifs discovered in the promoter regions of DEGs detected in *atpol2b* mutants using MEME Suite.

### Protein-protein interaction (PPI) analysis

Interestingly, the PPI network constructed for the DEGs identified in *atpol2a-1* using the StringDB showed that genes encoding REPLICATION PROTEIN A 1E (RPA1E; AT4G19130), TSO2 (AT3G27060) and AT3G49160 proteins may interact with AtPOL2A/TIL1 (Fig 9). Further, it was noted that RPA1E interacts with GAMMA-IRRADIATION AND MITOMYCIN C INDUCED 1 (GIM1, AT5G24280), and TSO2 interacts with POLY (ADP-RIBOSE) POLYMERASE 2 (PARP2; AT4G02390), and AT3G49160 interacts with FERRETIN 1 (FER1; AT5G01600).

**Fig 9.**
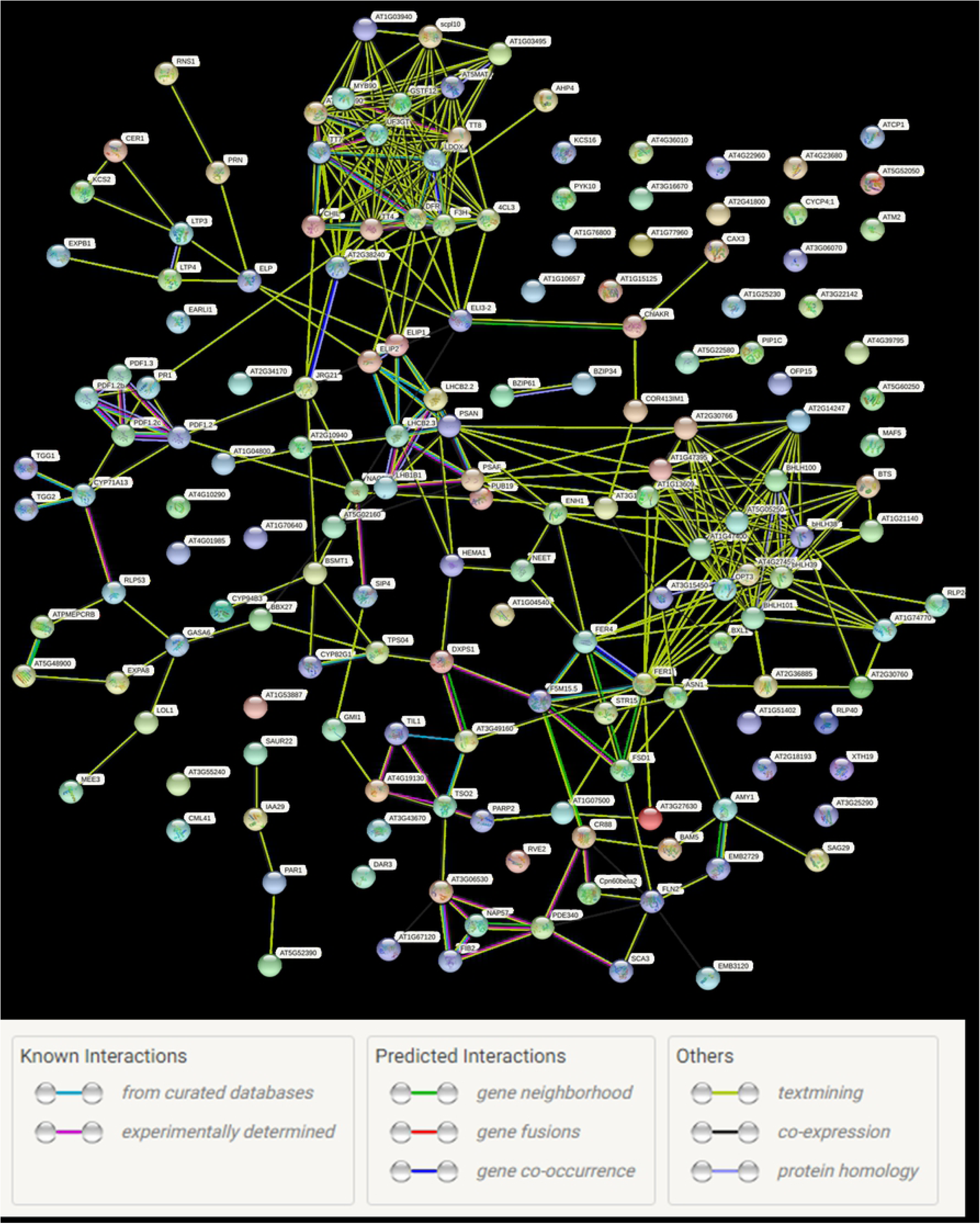
Protein-protein interaction network of DEGs detected in *atpol2a-1*. This network was constructed using StringDB. Network nodes represent proteins (colored nodes represent query proteins; empty nodes: proteins of unknown structures; filled nodes: some 3D structure is known or predicted.). Edges represent protein-protein interactions.

PPI network analysis of the DEGs detected in *atpol2b* mutants also showed known and predicted interactions among genes (Fig 10). Interestingly, of the 12 genes commonly differentially expressed in all three *atpol2b* mutants (see Table 3), genes encoding six proteins (KTI 1 (AT1G73260), PR1 (AT2G14610), PDF1.3 (AT2G26010), NIT2 (AT3G44300), OSM34 (AT4G11650) and ChiC (AT4G19810)) showed known/predicted interactions among them (Fig 10). However, none of the proteins were interacting with AtPOL2B/TIL2.

**Fig 10.**
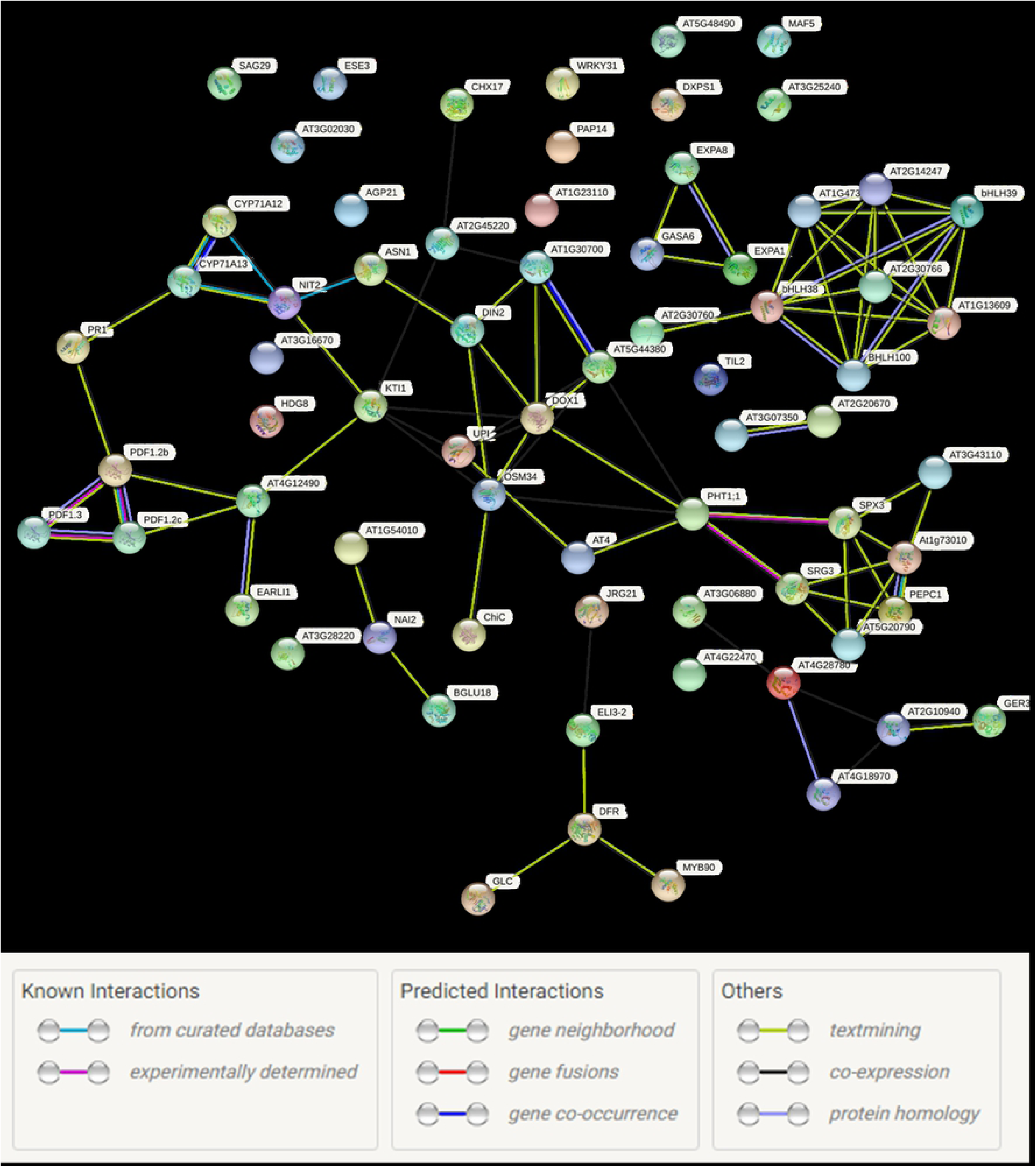
Protein-protein interaction network of DEGs detected in at least two *atpol2b* mutants. This network was constructed using StringDB. Network nodes represent proteins (colored nodes represent query proteins; empty nodes: proteins of unknown structures; filled nodes: some 3D structure is known or predicted.). Edges represent protein-protein interactions.

### Docking sites of potential targets of *AtPOL2*s

Analysis of the AtPOL2A and B interacting proteins through the StringDB showed that DNA polymerase epsilon subunit B2 (DPB2), DNA polymerase alpha subunit B (POLA2), DNA polymerase alpha primase subunit (POLA3), DNA replication licensing factor MCM2, DNA primase large subunit (EMB2813) and DNA polymerase delta small subunit (POLD2) are common interactors AtPOL2s (Fig 11). Docking scores obtained for these proteins are given in Table 5; the docking results demonstrated that the cluster size ranged from 38 to 152, while the balanced score ranged from -808.9 to -1174.5. Therefore, this scale was used to validate the interactions between AtPOL2s and several DEGs detected in the present study.

**Fig 11.**
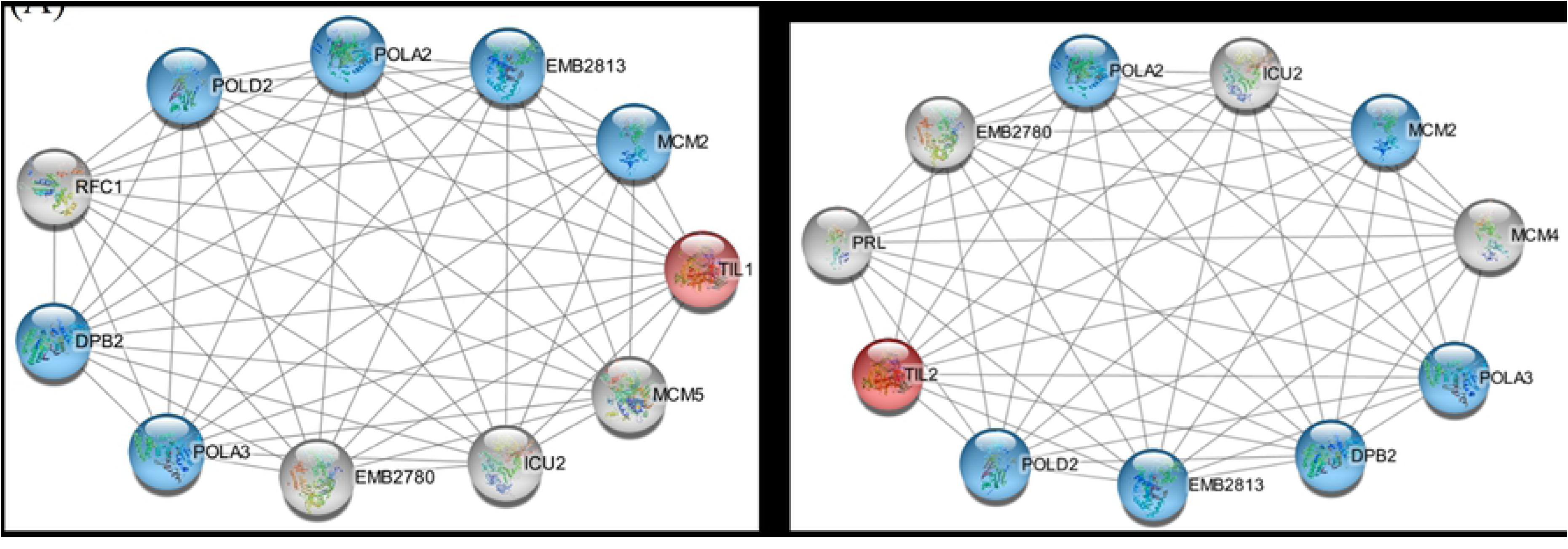
Protein-protein interactions of AtPOL2 proteins extracted from StringDB. The common interactors of AtPOL2A and AtPOL2B proteins are highlighted in blue. (A) Interactors of AtPOL2A (TIL1). (B) Interactors of AtPOL2B (TIL2).

**Table 5.**
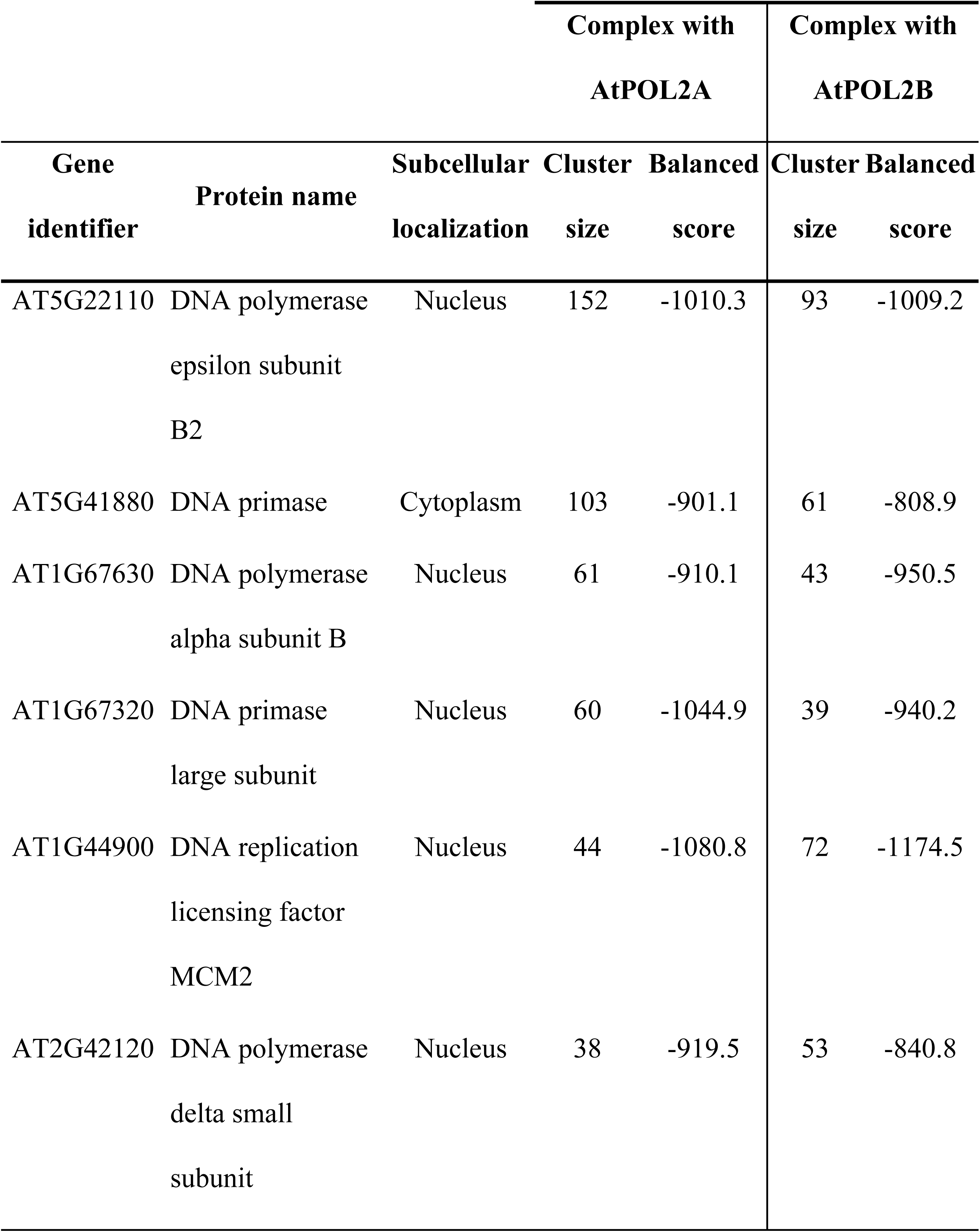
ClusPro docking scores for AtPOL2s and their known interactors.

Structural modeling of 22 DEGs (Table 1 and 3) using SWISS-MODEL predicted the 3D structures of 20 proteins, while two genes (AT3G27630 and AT2G14247) failed to locate homologous protein sequences for modeling. Out of the structures predicted, 90% (18) protein structures showed over 85% residues clustered in favoured regions in the Ramachandran plot, indicating stable conformations. Further analysis of these proteins revealed seven complexes with AtPOL2A and six complexes with AtPOL2B (*cluster size > 38* and *balanced score* less than -808.9; Table 6). Of these, three (CER1, RPA1E and AT5G60250) and two proteins (PR1 and AT5G48490) indicated notably significant interactions (*cluster size > 60* and *balanced score < -900*) with AtPOL2A and AtPOL2B proteins, respectively (Fig 12). Furthermore, these proteins also showed *cluster size > 60* and *balanced score < -900* when docked with the POLE catalytic modules (6HV8 and 6HV9) of *S. cerevisiae* (S2 Table).

**Fig 12.**
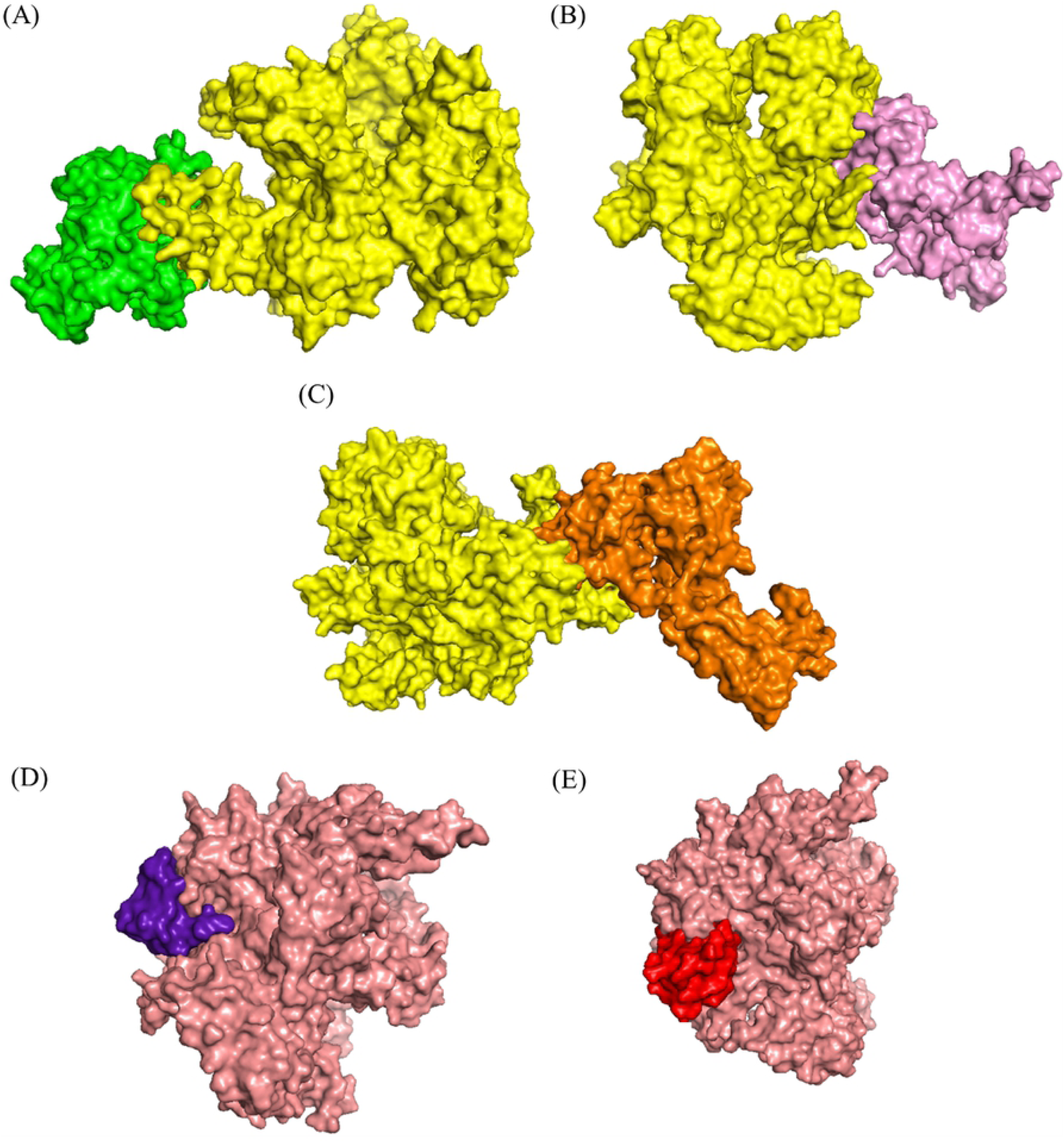
The 3-dimensional surface structures of potential AtPOL2A and AtPOL2B interactions. AtPOL2A and AtPOL2B are shown in yellow and salmon colours, respectively. The interaction sites of these protein docking complexes were visualized by PyMol. (A) AtPOL2A-CER1 interaction. (B) AtPOL2A-AT5G60250 interaction. (C) AtPOL2A-RPA1E interaction. (D) AtPOL2B-PR1 interaction. (E) AtPOL2A-AT5G48490 interaction.

**Table 6.**
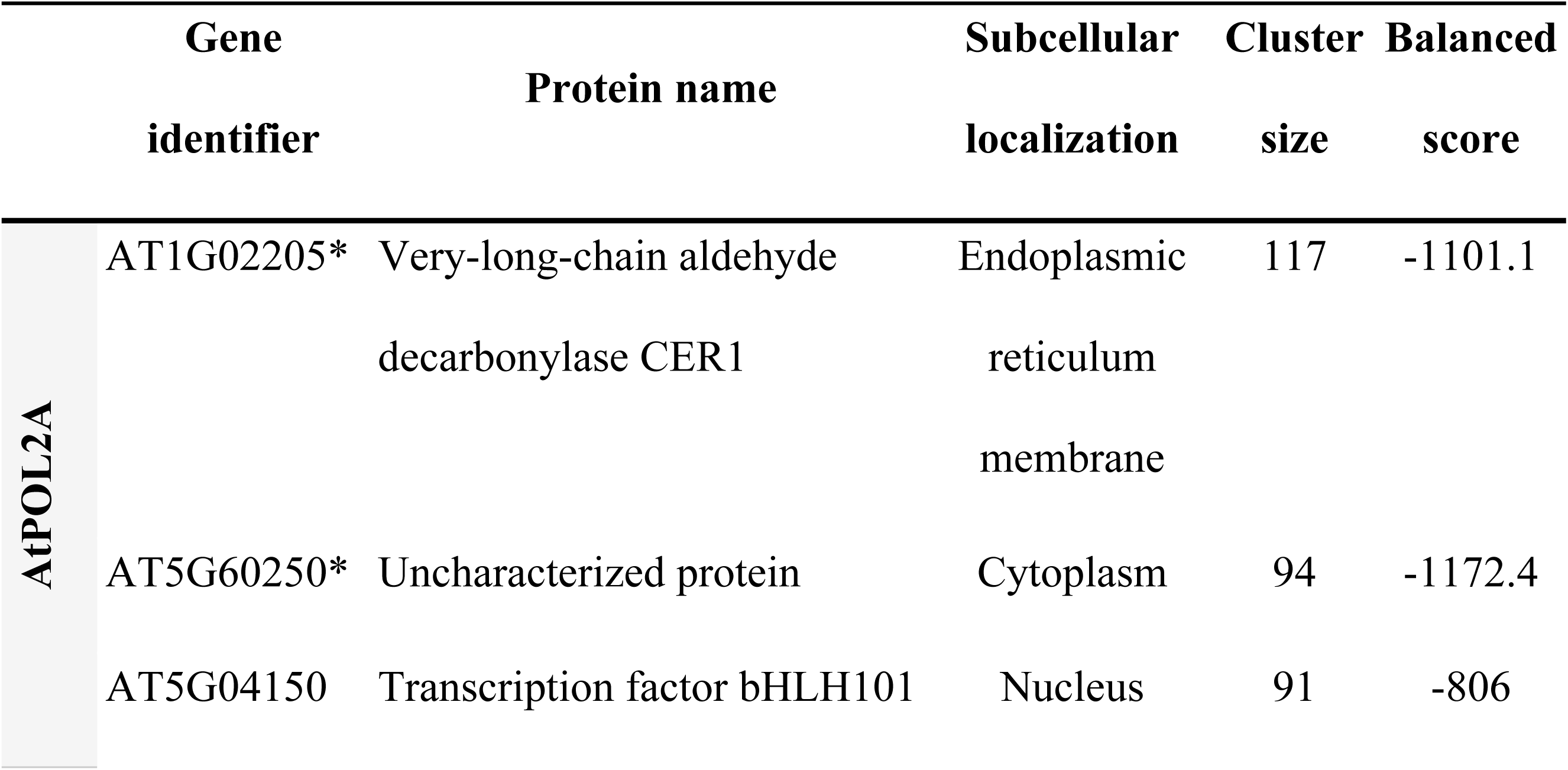
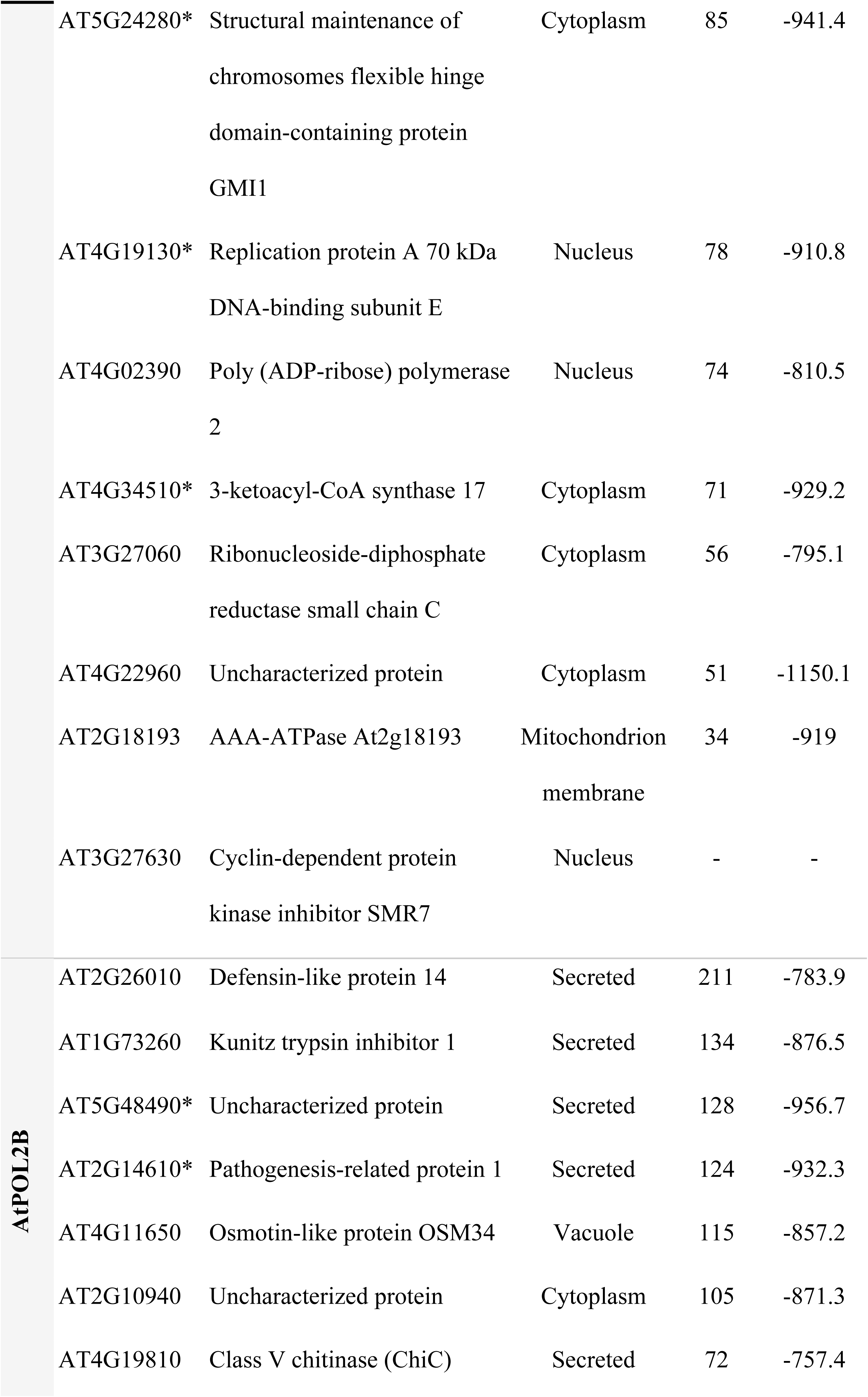
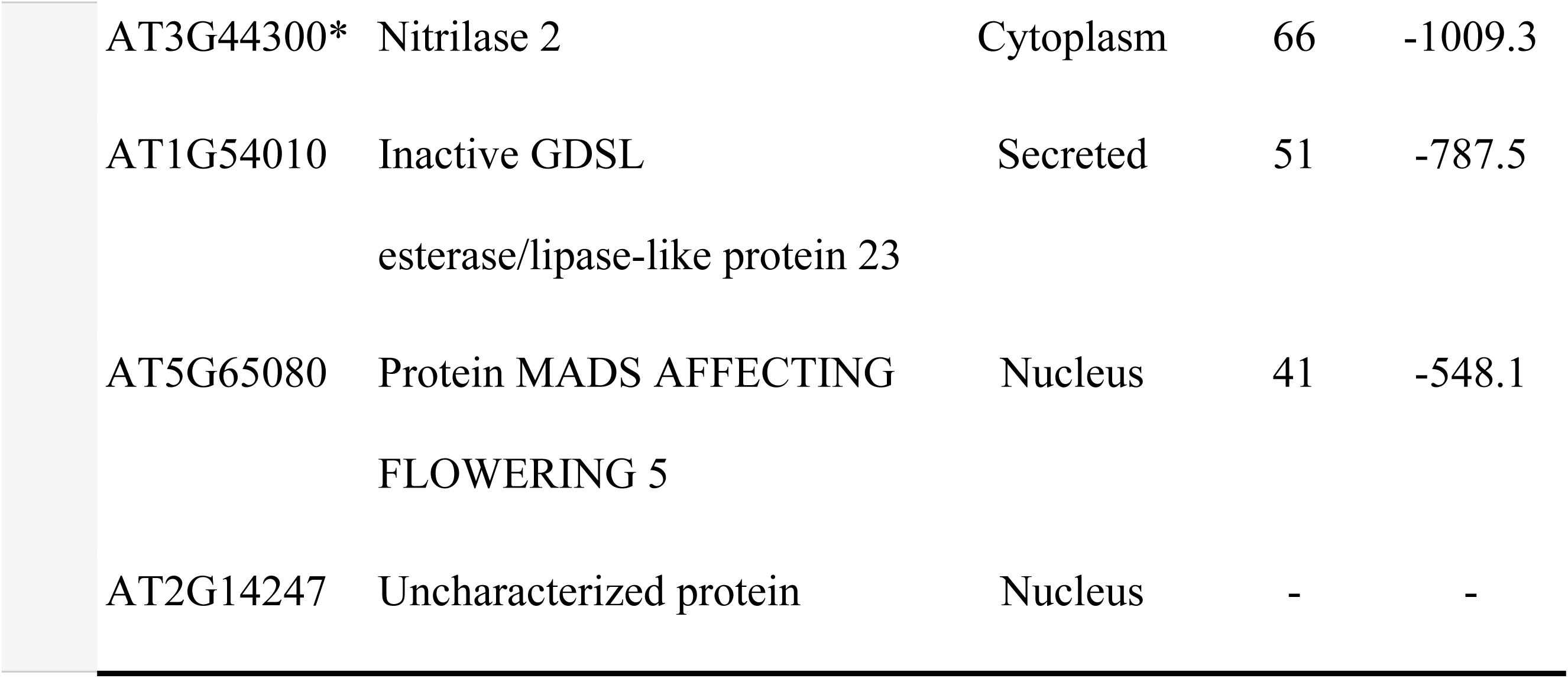
ClusPro docking scores for AtPOL2s and their potential targets. Significant protein interactions are highlighted with an asterisk (*cluster size > 60* and *balanced score < -900*).

### *In silico* analysis of *cis*-acting regulatory elements (CREs) of promoter sequences of *AtPOL2*s

The first insight into the promoter elements of *AtPOL2* genes came from a study conducted by Ronceret and co-workers in 2005 [9]. However, since then, no comprehensive analysis of the promoter sequences of these two genes has been performed. Therefore, an *in silico* promoter analysis was conducted to explore the potential interactors of *AtPOL2*s at the molecular level. The *AtPOL2A* gene contains a 2091 bp long 5’-untranslated region. Hence, 2500 bp upstream regions of both *AtPOL2* genes, from the translation start site (ATG) were selected and analysed using the PLACE and PlantCARE database to determine the putative CREs. The results obtained through the PLACE tool showed that a total of 113 and 110 putative CREs are distributed in *AtPOL2A* and *AtPOL2B* promoters, respectively; of these, 80 elements were found in both gene promoters. For instance, promoter motifs ARR1AT, DOFCOREZM, GATABOX, GT1CONSENSUS, GTGANTG10, MYCCONSENSUSAT, POLLEN1LELAT52, and WRKY71OS were most abundant in both gene promoters (Fig 13; descriptions of these CREs are included in S3A Table). In addition, CGCGBOXAT element (also known as ‘CGCG box’) involved in Ca^2+^/ calmodulin regulation, IBOXCORE element conserved in upstream of light-regulated genes, and MYBCORE element (consensus MYB recognition sequence) were overrepresented in the *AtPOL2A* promoter whereas CURECORECR (a copper-response element), INRNTPSADB (light-responsive initiator element) and RAV1AAT (recognition sequence of DNA-binding protein RAV1) were overrepresented in the *AtPOL2B* promoter.

**Fig 13.**
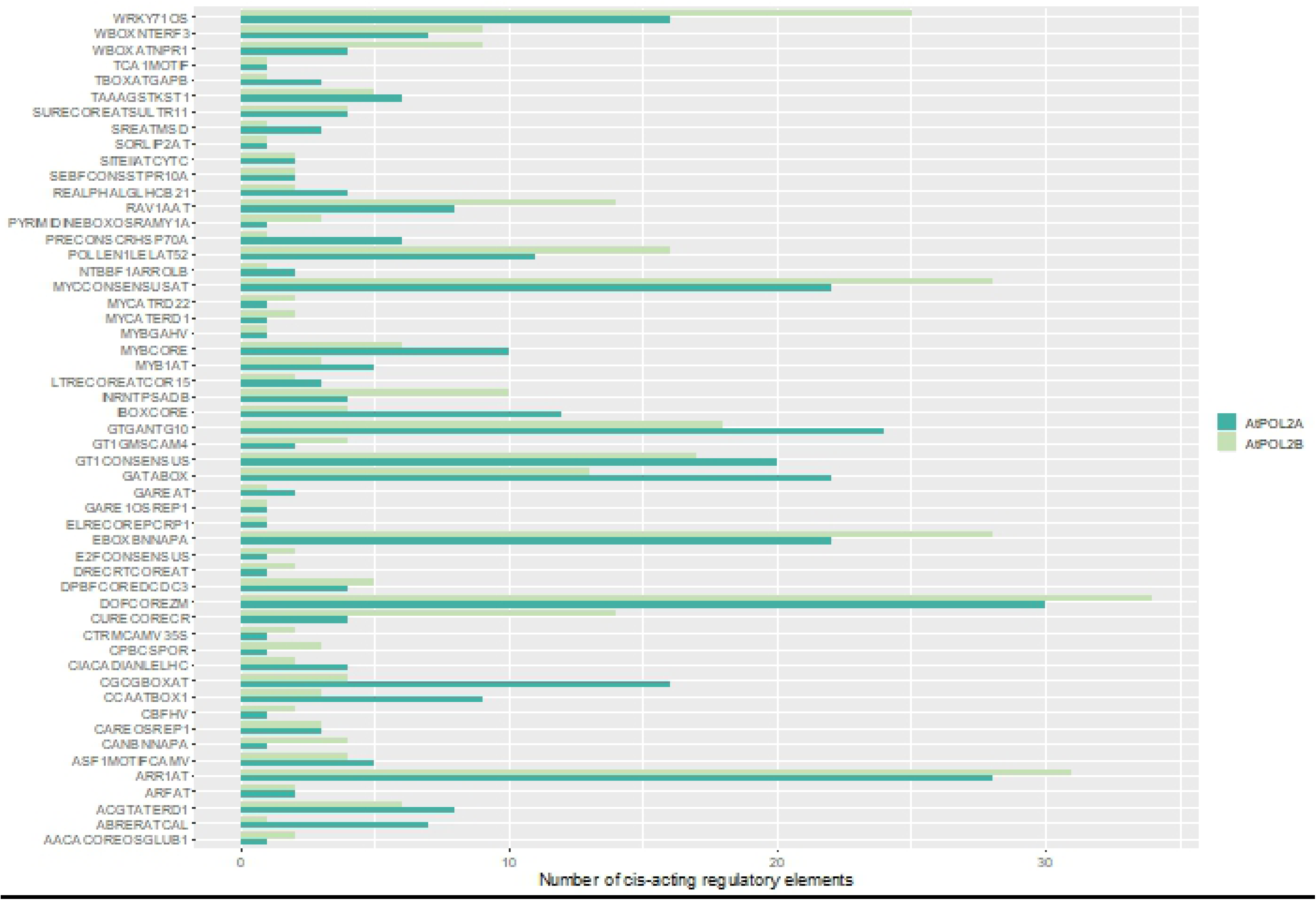
Frequencies of important CREs detected in promoter regions of both *AtPOL2* genes (2500 bp upstream from the translation start site)

Furthermore, the *AtPOL2A* gene promoter contained several CREs contributing to tissue-specific expression including pollen-specific elements (POLLEN2LELAT52 and VOZATVPP), embryo-specific elements (CARGCW8GAT), fruit-specific elements (TGTCACACMCUCUMISIN) and meristem-specific elements (E2F1OSPCNA). This gene also contained E2FANTRNR, an element related to up-regulation of the promoter at G1/S transition, and MYBCOREATCYCB1, a core CRE in *Cyclin B1; 1* gene in its promoter. In addition, several homeodomain-leucine zipper (HD-Zip) protein binding sites (ATHB2ATCONSENSUS and ATHB5ATCORE) and MYB-related binding sites (i.e. BOXLCOREDCPAL, CCA1ATLHCB1, IBOX, MYB2AT, MYB2CONSENSUSAT, MYBPLANT, MYBPZM) were detected.

Interestingly, a hexamer motif found in the Arabidopsis histone H4 gene promoter (HEXAMERATH4) was also detected in the *AtPOL2A* promoter. By contact, QELEMENTZMZM13 (Quantitative elements (Q-element)), which is known to enhance activity in pollen-specific genes; AUXRETGA1GMGH3, an auxin-responsive CRE; LTRE1HVBLT49, a low-temperature responsive element; PREATPRODH, a hypoosmolarity-responsive element; DRE2COREZMRAB17, SBOXATRBCS and MYBATRD22 (drought and/or ABA-responsive elements); SORLIP5AT (a light-responsive element); HEXMOTIFTAH3H4, a conserved hexamer motif known to interact with histone DNA binding proteins; WBBOXPCWRKY1 and WBOXNTCHN48 (pathogen/elicitor responsive elements) were detected only in the *AtPOL2B* promoter. Important CREs detected only in one of the *AtPOL2*s gene promoters and their functions are listed in S3B and S3C Tables.

Analysis of the promoter sequences using PlantCARE also showed that both *AtPOL2* promoters contained hormone-responsive elements such as auxin-responsive, gibberellin responsive and methyl jasmonate responsive; however, ABA-responsive and salicylic acid responsive elements were found only in the *AtPOL2A* promoter sequence (Fig 14A). Among the environmental stress-responsive elements, drought-responsive elements were only detected in the *AtPOL2A* promoter, and defense-responsive and low-temperature responsive elements were only detected in the *AtPOL2B* promoter (Fig 14B). Further, it was noted that both *AtPOL2* genes contained a considerable number of light-responsive elements. As expected, an element responsible for the regulation of the cell cycle was only detected in the *AtPOL2A* promoter sequence (Fig 14C).

**Fig 14.**
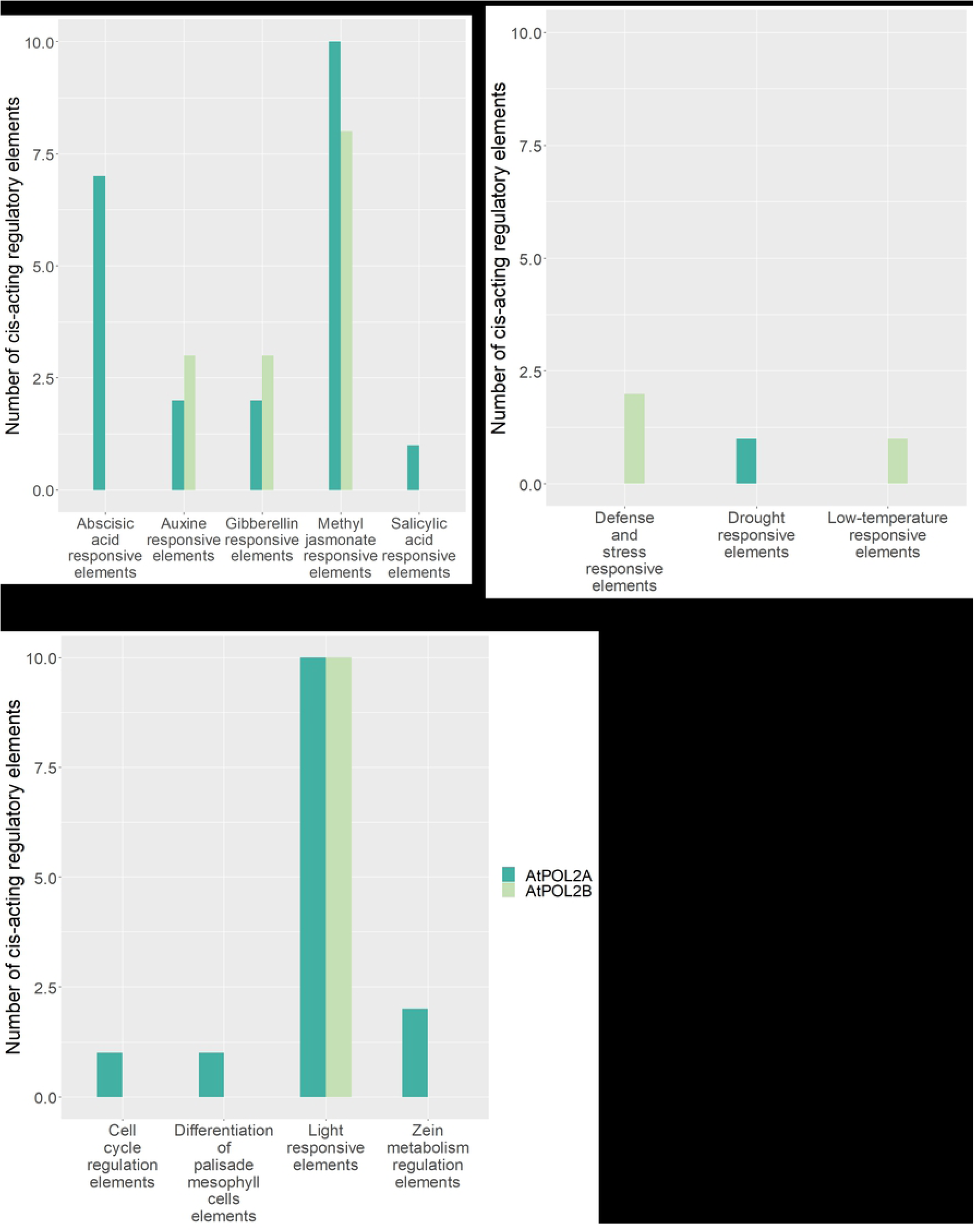
Abundance of CREs identified using PlantCARE in the 2500 bp upstream region of each *AtPOL2* gene. (A) Phytohormone responsive CREs. (B) Environmental stress responsive CREs. (C) Plant growth and developmental processes related CREs.

## Discussion

DNA Pols play a central role in maintaining genomic integrity. In the present study, we investigated the potential indirect targets of the two isoforms of the catalytic subunit of Arabidopsis POLE, AtPOL2A and AtPOL2B at the transcriptomic level. We further identified the putative CREs in the promoter regions of *AtPOL2* genes to better understand the potential targets of *AtPOL2* genes.

### Transcriptome analysis of *AtPOL2* gene mutants

The publicly available T-DNA insertional mutant line collections have played an important role in functional studies to explore gene functions and gene/protein regulatory networks. In the present study, we employed four homozygous T-DNA insertional lines. All of these mutants, except *atpol2b-3*, contained insertions in introns. However, all mutants showed a change in the expression pattern of the mutated gene as compared to that of in the WT; although Ronceret *et al*. (2005) report no expression of *AtPOL2B* in *atpol2b* mutants (SALK_001413 and SALK_056503), we detected increased transcript accumulation downstream of the insertion site in all *atpol2b* mutants. Detection of transcripts both upstream and downstream of the T-DNA insertion site is supported by previous literature [18]. Such fusion/truncated transcripts formed due to activation of T-DNA promoter or termination of transcription due to inserted T-DNA fragment could translate either into a functional or a truncated protein [19]. A closer look at the primer sequences used by Ronceret and co-workers [13] to amplify *AtPOL2B* transcripts showed that they have been designed from regions upstream of the T-DNA insertion site. This might be the most likely reason for the failure to detect *AtPOL2B* expression in *atpol2b* mutants in the previous literature.

Consistent with previous findings [14], the *AtPOL2A* mutant examined in the present study (*atpol2a-1*) was also early flowering and exhibited smaller siliques than WT/*AtPOL2B* mutants. This observation, together with the expression pattern of the mutated gene in the T-DNA insertional line, strongly suggests *atpol2a-1* is a weak allele of *AtPOL2A*. Moreover, as reported in previous studies [4,13,15], none of the homozygous *AtPOL2B* mutant lines examined here exhibited visible phenotypic variation as compared to the WT. However, as all of them showed a considerable change in the expression pattern of the mutated gene, they were used in subsequent transcriptomic analysis to explore potential downstream targets of *AtPOL2B*.

Several different bioinformatics tools have been developed and used in the analysis of expression data generated through next-generation technologies. However, having multiple tools for analysis has made the analysis and interpretation of data challenging for researchers and has led to large variations in outputs [20, 21]. Hence, in the present study, two computational tools for transcript assembly (Cufflinks and HTSeq) and three tools for differential gene expression analysis (cuffdiff, edgeR and DESeq) were used; only DEGs resulted from all the tools were considered for the downstream analysis.

This resulted in a total of 198 genes in *atpol2a-1*. Han *et al*. (2015) [17] and Pedroza-Garcia *et al*. (2017) [16] report 749 ( -1 ≥ Log_2_ (FC) > 1; P < 0.01) and 372 ( -1 ≥ Log_2_ (FC) ≥ 1.5; P ≤ 0.01) DEGs in a single-gene mutant of *AtPOL2A*, respectively. The discrepancies observed in DEGs between mutants of *AtPOL2A* are likely to result from the differences in the plant material and growth conditions, sequencing platforms used to generate transcriptomic data, sequencing depth, number of replicates and analysis methods. Despite discrepancies, we identified 11 DEGs in all *AtPOL2A*-deficient mutants. This included several genes involved in the DNA damage and repair response pathways such as *TSO2*, *SMR7*, *RPA1E*, *PARP1* and *GMI1*. Further analysis of these DEGs through PPI network analysis showed that TSO2, RPA1E and AT3G49160 (a pyruvate kinase-related protein) [22] may associate with AtPOL2A; this was further supported by the protein-protein docking using Cluspro. In Arabidopsis, *TSO2* is one of the three members of the ribonucleotide reductase (RNR) small subunit genes, which are transcribed predominantly at the S-phase of the cell cycle [23]. Roa *et al*. (2009) [24] report E2F target genes including *AtPOL2A* and *TSO2* coexpress in the DNA repair network. The *tso2* mutants have also shown increased DNA damage, the release of transcriptional gene silencing and sensitivity to UV-C light as observed in *atpol2a* mutants [23]. Moreover, *tso2-1 rnr21-1* double mutants have shown induced expression of several molecular markers associated with DNA damage and repair i.e. *PARP1*, *PARP2* and *RAD51* [23]. Recently, upregulation of *PARP2*, *RPA1E* and *POL2A* have been observed in loss-of-function mutants of the F-box protein *F-BOX-LIKE17* (*FBL17*); in Arabidopsis, *FBL17* functions in cell-cycle regulation [25], and is required for cell division during pollen development. Our PPI analysis also showed that uncharacterized AT3G49160 interacts with FER1; both of these genes are known to be expressed in response to hydrogen peroxide [26–28].

Mutations in several genes related to DNA/histone assembly and replication have been reported to release the transcriptional gene silencing of pericentromeric *Athila* retrotransposon at the *transcriptional silent information* (*TSI*) i.e. *abo4-1* and *abo4-2* mutants [14, 17]. TEs found in the pericentromeric heterochromatin are heavily methylated. In a recent study, Bourguet and his colleagues have noted that the significant release of heterochromatin silencing in *AtPOL2A* gene mutants is due to increased levels of DNA methylation and histone H3 lysine 9 methylation, and thus *AtPOL2A* is important to maintain heterochromatin structure and function [29]. Consistent with previous studies, we also observed that the silencing of the majority of pericentromeric TEs are released in *atpol2a-1* mutant.

Notably, the analysis of the expression profiles of three *atpol2b* mutants showed that the DEGs in mutants are random; remarkably, there was little overlap between the *atpol2b* mutant-derived DEGs. It is reported that POLE may be involved in the repair of replication errors [16]; the *POLE* mutants might not be able to repair errors during DNA replication and such replication errors can occur randomly. Therefore, in different pole mutants, the DEGs could be random.

Interestingly, our transcriptome analysis showed that *AtPOL2B* is more likely to interact with stress-responsive genes; several genes related to biotic stress responses were differentially expressed in all three *atpol2b* mutants examined i.e. *GLL23*, *KTI1*, *PR1* (a salicylic acid marker gene) [30, 31], *PDF1.3* (a plant defensin) [32], *OSM34, DIR1-LIKE* (may contribute to systemic acquired resistance) [33] and *ChiC*. Phylogenetic analysis of GDSL-type esterases/lipases has identified four clades (I –IV), and *GLL 23* has been placed in clade IIIa; T-DNA knockout lines of many genes in this clade have shown increased resistance to biotic stresses [34]. Moreover, *KTI1* (serine protease inhibitor) is known to be involved in the regulation of programmed cell death during biotic stress tolerance in Arabidopsis [35, 36]. In addition to defense-related processes, *OSM34* has shown to be effective against abiotic stresses [37–39]. Furthermore, the *ChiC* gene which was also differentially expressed in all *AtPOL2B* mutants is known to be induced by the plant stress-related hormones (ABA, jasmonic acid) and by the stress resulting from the elicitor flagellin, NaCl and osmosis [40]. In addition to these genes, a member of a NITRILASE 1-subfamily, *NIT2* was also differentially expressed in all *atpol2b* mutants. *NIT2* regulates hypocotyl elongation in response to high temperatures [41]. Interestingly, protein-protein docking of these DEGs confirmed significant interaction between AtPOL2B and NIT2, OSM34 and Chic proteins.

Under our experimental conditions, *MAF5* was differentially expressed in both *atpol2a* and *b* mutants compared to WT. This finding is contradictory to the results obtained by del Olmo *et al*. (2010); no significant alteration in floral development genes including *MAF5* has been observed in *AtPOL2A* mutant *esd7-1* [15]. However, Han *et al.* (2015) also report differential expression of *MAF* in the *AtPOL2A* mutant, *abo4-1* [17]. Previous studies have shown that early flowering mutants of Arabidopsis exhibit a reduction in levels of H3 methylation (H3K4me3 and H3K36me2) at the flowering locus genes *FLC*, *MAF1*, *MAF4* and *MAF5* [42, 43]. These observations suggest *AtPOL2A* is likely to regulate the methylation status of *MAF5* and thereby control the expression of *MAF5*.

### Computational analysis of *AtPOL2* promoter regions

As reported in Ronceret *et al*., 2005 [13], one E2FANTRNR element (also known as E2Fa element), which is a binding site for E2F-like proteins, was detected upstream of the translation start site of *AtPOL2A*. In addition, E2F1OSPCNA, which is also an E2F transcription factor binding site, was identified in the promoter region of *AtPOL2A*. In Arabidopsis, consensus E2F-binding sites have been identified in the promoter regions of several regulators of the cell cycle (e.g. D-type cyclin CycD3 and CDC6) suggesting a potential role of E2F and E2F-like proteins in G1-to-S phase transition of the cell cycle [44]. Furthermore, Kosugi and Ohashi (2002) report, E2F binding sites are required for meristematic tissue-specific activity of proliferating cell nuclear antigen promoters (PCNA) in rice and tobacco [45]. Neither E2FANTRNR nor E2F1OSPCNA elements were found upstream of the translation start site of *AtPOL2B*. However, E2FCONSENSUS (an E2F consensus sequence) and Myb-binding core (AACGG) motifs were detected in both promoter regions; Myb-binding core, AACGG is responsible for cell cycle phase-independent transcriptional activation of B-cyclin *CycB1;1* in Arabidopsis [46]. Analysis of the microarray data has further confirmed that Arabidopsis E2F target genes are expressed during G1 and S phases and most of the proteins encoded by these genes function in cell cycle regulation, DNA replication and chromatin dynamics [47].

Moreover, both *AtPOL2A* and *B* promoters contained motifs reported to be conserved in the promoter regions of plant and animal histone genes; HEXAMERATH4, a hexamer motif (CCGTCG) found in histone H4 promotes [48], and HEXMOTIFTAH3H4, a hexamer motif (ACGTCA) found in H3 and H4 histone gene promoters [49] were found in *AtPOL2A* and *AtPOL2B* promoters, respectively. The hexamer motif CCGTCG found in the wheat histone genes has been shown to be essential for a meristematic-tissue specific pattern of gene expression i.e. expression in organs containing meristematic tissues such as roots and flower buds than in fully expanded leaves and therefore, promoters having CCGTCG motifs could exhibit a cell-division inducible activity [48]. Moreover, the conserved hexamer motif ACGTCA found in the plant histone H3 and H4 genes binds specifically with the nuclear protein HBP-1 (histone gene-binding protein-1) [50–53], which is more abundant in actively proliferating cells; it is reported that HBP-1 may be involved in cell-cycle dependent expression of histone genes in plants. Detection of an ACGTCA motif upstream of the translation initiation codon of *AtPOL2B* hence suggests cell-cycle dependent transcription of *AtPOL2B*.

Notably, DOFCOREZM and GTGANTG10 were abundantly found in both *AtPOL2* promoter sequences. DOFCOREZM elements specifically recognize plant-specific Dof (DNA-binding with one finger) TFs, which act as transcriptional regulators involved in many plant-specific biological processes [54–56]. Engel *et al*. (2005) [57] report the presence of core binding site for Dof transcription factors in promoter sequences of two sperm-specific genes *AtGEX1* (AT5G55490) and *AtGEX2* (AT5G49150). Furthermore, GTGANTG10 is a GATA motif found within the promoter of the tobacco late pollen gene g10; Rogers *et al*. (2001) have demonstrated that this gene is expressed preferentially and maximally in mature pollen [58].

Interestingly, two co-dependent CREs, POLLEN1LELAT52 (AGAAA) and POLLEN2LELAT52 (TCCACCATA) were found in the *AtPOL2A* promoter. Both of these CREs are required for the tissue-specific regulation of late pollen genes during pollen development [59]. Also, two VOS-binding motifs were found upstream of the translation initiation codon of *AtPOL2A*; Arabidopsis VOS proteins (AtVOZ1 and AtVOZ2) function as transcriptional activators [60]. The pollen-specific transcriptional activation of the Arabidopsis *AVP1* gene that encodes vacuolar H+-pyrophosphatase (V-PPase) is controlled by binding of AtVOZ1 and AtVOZ2 transcription factors to a 38 bp pollen-specific CRE [61]. Moreover, two CWWWWWWWWG motifs (a variant of CArG motif) that interact with AGAMOUS-like 15 (AGL15) were found in the *AtPOL2A* promoter. The MADS domain family transcription factor AGL15 preferentially accumulates during embryo development [62]. Further, a TGTCACACMCUCUMISIN (TGTCACA) was found in the promoter region of *AtPOL2A*; this enhancer element is responsible for the fruit-specific expression of cucumisin gene in melon (*Cucumis melo* L.) [63]. Detection of AGL15 binding sites and TGTCACACMCUCUMISIN motif in the *AtPOL2A* gene further confirms the function of *AtPOL2A* in Arabidopsis embryo development. By contrast, three QELEMENTZMZM13 (Q-element) and 16 POLLEN1LELAT52 motifs were identified within the regulatory region of *AtPOL2B*. The Q-element has been identified in a promoter region of a pollen-specific gene (*ZM13*) in maize [64]. However, the pollen-specific expression of *ZM13* is known to be regulated by two motifs (pollen-specific and Q element) in the promoter; the presence of only the Q-element in the promoter has no ability to cause the expression of *ZM13* in pollen [64]. These findings further confirm *AtPOL2B* is more likely to exhibit no or little expression in reproductive tissues.

The present study also found two binding sites for Arabidopsis HOMOMEOBOX 2 (AtHB2) and two binding sites for AtHB5 in the *AtPOL2A* promoter. Of these HD-Zip class proteins, *AtHB2*, which expresses during both vegetative and reproductive stages of plant growth, is found to be strongly induced in response to Red to Far-Red light ratio [65, 66], whereas *AtHB5* appears to be involved in ABA-related responses and/or water deficit conditions [67]. In addition, several MYB protein recognition sequences were found in the promoter region of *AtPOL2A* (BOXLCOREDCPAL, CCA1ATLHCB1 [68], MYB2CONSENSUSAT, MYBPLANT (a binding site for a flower-specific MYB protein [69]), MYBPZM, MYB2AT and IBOX [70]). In plants, including hormonal signal transduction and abiotic/biotic stress tolerance [71, 72], and transition [73].

We also noted that several light-responsive elements (GATABOX) and stress responsive elements (MYCCONSENSUSAT, ARR-1 binding elements [74, 75], WRKY71OS [76] are distributed within both *AtPOL2* promoters. However, as compared to the *AtPOL2A* promoter sequence, many stress-responsive elements were found in the *AtPOL2B* promoter. i.e. abiotic stress responsive elements (LTRE1HVBLT49, PREATPRODH, DRE2COREZMRAB17, SBOXATRBCS and MYBATRD22) and biotic stress responsive (pathogen/elicitor responsive) elements (WBBOXPCWRKY1 and WBOXNTCHN48). Transcriptome analysis of *atpol2b* mutants together with the promoter analysis confirm *AtPOL2B* is more likely to play a role in modulating stress responses in Arabidopsis.

## Conclusion

This study, based on several *AtPOL2* mutants, is the first to present a detailed comparison of the transcriptomes of *AtPOL2A* and *AtPOL2B* mutants on a global scale. Most importantly, the findings of the present study begin to unravel the potential targets of *AtPOL2* genes at the molecular level and suggest new avenues for future studies.

## Materials and Methods

### Planting materials and growth conditions

Seeds of *A. thaliana* (ecotype Columbia-0) T-DNA insertion lines of *AtPOL2A* [SALK_096314C (*atpol2a-1*)] and *AtPOL2B* [N859810 (*atpol2b-1*), SALK_001413 (*atpol2b-2*), SALK_056503 (*atpol2b-3*)]; WT (N70000) were obtained from the Nottingham *Arabidopsis* Stock Centre, UK. Seeds were germinated and grown under controlled environmental conditions at 25°C, 60% relative humidity and 16 h photoperiod (Fitotron plant growth chambers, Weiss Gallenkamp, UK).

### Morphological characterization

The average silique length and the number of days to flower were recorded for mutants and WT plants. Green siliques were harvested 12 d after flowering from 4-6 week old plants and opened using fine needles under a dissecting microscope (Leica MZ9.5, Germany). Images of opened siliques were captured using a Leica DFC290 (Germany) camera and edited in Adobe Photoshop.

### Molecular screening of T-DNA mutants

Genomic DNA was extracted from leaves of mutants and WT plants using the DNeasy^®^ Plant Mini Kit (Qiagen, UK) according to the manufacturer’s instructions. Homozygous mutant lines were identified through a PCR-based screening method using the primers designed through the SIGnAL iSct tool (http://signal.salk.edu/tdnaprimers.2.html). The list of primers used to confirm T-DNA mutant lines used in the present study is given in S4 Table.

In brief, two main PCR reactions with primer combinations, (1) LB and RB genomic primers; (2) LB/RB primer and a T-DNA specific primer (LBb1.3 or LBa1) were set up to confirm mutant lines. Amplifications were performed in a volume of 25.0 µl, containing 12.5 µl of 2X BioMix PCR master mix (Bioline, UK), 0.75 µl of 0.3 µM forward and reverse primer (Invitrogen, UK) and 1.0 µl of genomic DNA. Each amplification was carried out in a GeneAmp PCR system 9700 (Applied Biosystems) using the following programme: 2 min at 94°C; followed by 30 cycles of 30 s at 94°C, 30 s at 55°C, and 1 min at 72°C; and finally 10 min at 72°C for the final elongation. All the amplified products were confirmed by sequencing at a commercial company (Source Bioscience, UK).

### Preparation of cDNA libraries and whole transcriptome sequencing

Two fully expanded leaves were harvested from each confirmed homozygous mutant line and WT plants at the time of development of the first flower bud. Total RNA was extracted using the RNeasy^®^ Plant Mini Kit (Qiagen, UK) according to the manufacturer’s instructions and quantified through Agilent BioAnalyzer 2100. cDNA libraries were constructed and amplified using random primers according to the manufacturer’s instructions (Illumina TruSeq RNA sample preparation guide v2 for Illumina paired-end multiplexed sequencing). For sequencing, two samples were pooled and loaded into each flow cell and sequenced using Illumina HiSeq 2000 platform (2 x 50 bp read length).

### Aligning sequence reads to the *A. thaliana* reference genome

The quality of raw sequence reads generated through sequencing was assessed using the FastQC tool (http://www.bioinformatics.babraham.ac.uk/projects/fastqc/). After clearing for adapter sequence contaminations, TopHat 2.1.1 [77] (http://tophat.cbcb.umd.edu) with Bowtie2 2.3.4.1 (http://bowtie-bio.sourceforge.net/index.shtml) was used to align sequences to the Arabidopsis reference genome (TAIR10), allowing only 2 base mismatches per read alignment [78]. The Cufflinks (2.1.1) was used to assemble reads that were mapped and to normalize transcript levels [79] by means of Fragments Per Kilobase of exon per Million fragments mapped (FPKM) [79]; in addition, HTSeq-count 0.11.1 [80] (https://htseq.readthedocs.io/en/release_0.11.1/count.html) program with the intersection ‘union’ option was also employed to estimate gene-level expression of read alignments sorted by ‘name’ using SAMtools 1.11 (http://www.htslib.org/doc/1.11/samtools.html).

### Identification of DEGs and downstream analysis

Three tools based on the negative binomial distribution, cuffdiff (a part of the Cufflinks package) [81], DESeq [82] (https://bioconductor.org/packages/release/bioc/html/DESeq.html) and edgeR [83] (https://bioconductor.org/packages/release/bioc/html/edgeR.html) were used for the differential gene expression analysis; of these DESeq and edgeR can run on the dataset without replicates, and cuffdiff can be used to compare the log-ratio of gene expression (FPKM) between mutants. FC ratio of transcripts between samples was used to filter DEGs; the genes that fell within -1 ≥ Log_2_ FC ≥ 1 with an adjusted P-value of ≤ 0.05 were considered as differentially expressed. Functional enrichment analyses were performed using the DAVID v6.8 [84]. GO terms and KEGG pathways were considered enriched if the associated Benjamini-Hochberg adjusted p-value was less than 0.05.

The promoter sequences (1000 bp upstream of TSS) of each DEG were retrieved from the TAIR (https://www.arabidopsis.org/). The motif-based sequence analysis tool, MEME Suite (https://meme-suite.org/meme/tools/meme) was used to discover overrepresented motifs in DEGs; biological roles of predicted motifs were determined using the GOMo (Gene Ontology for Motifs) tool (https://meme-suite.org/meme/tools/gomo).

### Interactome analysis and docking sites of potential targets of AtPOL2s

In order to identify proteins that interact with AtPOL2A and B, an interactome analysis was done using STRING database (https://string-db.org/). All proteins encoding genes that were differentially expressed in AtPOL2A/AtPOL2B mutants were used as the set of potential interactors and default parameters were used to construct the network (interaction score > 0.400).

The protein sequences of the 22 DEGs (Table 1 and 3) were retrieved from the TAIR database using ‘Bulk data retrieval’ tool. The 3D structures of the proteins were generated using SWISS-MODEL (https://swissmodel.expasy.org/). Each predicted structure was confirmed using Ramachandran plots using the MOLProbity program [85]. In addition, the subcellular localization of the proteins was explored using the LocTree3 tool (https://rostlab.org/services/loctree3/). The ClusPro server 2.0 (https://cluspro.bu.edu/), which employs a rigid body protein-protein docking [86], was used to predict the docking sites for the AtPOL2 proteins with their target proteins. The best docking structure was selected based on the largest cluster size and minimum balanced score and visualized through PyMOL molecular graphics system (version 2.0). In addition, proteins interacting with AtPOL2A and AtPOL2B were identified using StringDB, structurally modeled through SWISS-MODEL. Further, the existing 3D structures of POL2 proteins in *S. cerevisiae* were retrieved from PDB (PDB accession numbers: 6HV8 and 6HV9; https://www.rcsb.org/). These protein structures were docked with the predicted structures of the putative tragets of AtPOL2 using ClusPro 2.0.

### Confirmation of *AtPOL2* expression through RT-qPCR

Total RNA was extracted from leaf tissues of mutant and WT plants using the RNeasy^®^ Plant Mini Kit (Qiagen, UK) according to the manufacturer’s instructions. First-strand cDNA was synthesized using SuperScript III First-Strand Synthesis SuperMix for qRT-PCR (Invitrogen, UK) using 500 ng of total RNA.

Gene sequences of *AtPOL2A* (*AT1G08260*) and *AtPOL2B* (*AT2G27120*) and *Arabidopsis UBC21* (*AT5G25760*) were retrieved from GenBank (http://www.ncbi.nlm.nih.gov). Primers were designed using the NCBI-Primer BLAST tool (http://www.ncbi.nlm.nih.gov/tools/primer-blast). One primer of each pair was designed, where possible, to span an exon-exon junction (see S5 Table).

RT-qPCR was performed in 20.0 µl final reaction volume containing 10.0 µl of 2X SensiMix SYBR^®^ No-ROX master mix (Bioline, UK), 0.3 µM of forward and reverse primers and 1.0 µl of cDNA. PCR was carried out on a Rotor-Gene 6000 System (Corbett Life Sciences, UK). Thermal cycling conditions consisted of one cycle of 95°C for 10 min; then 40 cycles of denaturing at 95°C for 15 s and annealing/elongation at 60°C for 1 min, followed by melting curve analysis from 60 to 95°C at the rate of 0.5°C per s. Each sample was repeated thrice. The fluorescence was recorded during the annealing/elongation step in each cycle. Primer specificity was verified by the melting curve analysis, and by DNA sequencing of the PCR products. The relative standard curve method was employed for gene expression analysis. Five-fold serial dilutions of cDNA were used to construct standard curves. The slope of the standard curve was used to estimate the PCR amplification efficiency. The expression of the target *AtPOL2* genes was normalized to the *UBC* reference gene. Data were obtained and analyzed using Rotor-Gene 6000 series software Version 1.7.

### *In silico* promoter analysis of *AtPOL2*s

Promoter sequences (2500 bp upstream of the translation start site) of each *AtPOL2* gene were retrieved from the TAIR (https://www.arabidopsis.org/). PLACE (database of plant c*is*-acting regulatory DNA elements; http://www.dna.affrc.go.jp/PLACE/) and PlantCARE databases were employed for screening of putative CREs (http://bioinformatics.psb.ugent.be/webtools/plantcare/html/). Additionally, novel motifs were discovered using MEME.

## Abbreviations

ABA: Abscisic acid
ABO: ABSCISIC ACID OVERSENSITIVE
ChiC: Class V chitinase
CRE: *Cis*-acting regulatory element
DEG: Differentially expressed gene
DNA: Pols DNA polymerases
ESD: EARLY IN SHORT DAYS
GO: Gene Ontology
HBP-1: Histone gene-binding protein-1
LB: Left border
NIT2: NITRILASE 2
POLE: DNA polymerase ε
PPI: Protein-protein interaction
PR1: PATHOGENESIS-RELATED GENE 1
Q-element: Quantitative element
RB: Right border
RPA1E: REPLICATION PROTEIN A 1E
TE: Transposable element
TIL: TILTED
TSS: Transcription start site
WT: Wild type

## Acknowledgments

Not applicable.

## Supporting information

**S1 Figure. Molecular confirmation of the *atpol2a-1* mutant.**

**S2 Figure. Molecular confirmation of *atpol2b* mutants.**

**S1 Table List of DEGs detected in *atpol2a* and *atpol2b* mutants as compared to the WT.** This file lists the DEGs detected in *atpol2a* (S1A Table), *atpol2b-1* (S1B Table), *atpol2b-2* (S1C Table) and *atpol2b-3* (S1D Table) mutants as compared to the WT.

**S2 Table. ClusPro docking scores for DEGs in *atpol2a* and *atpol2b* mutants with POL2 protein of *S. cerevisiae*.**

**S3 Table. List of important CREs detected in genomic regions (2500 bp upstream of the translation start site) of *AtPOL2A* and *AtPOL2B*.** This file lists the CREs detected in both *AtPOL2A* and *AtPOL2B* (S3A Table), only in *AtPOL2A* (S3B Table) and only in *AtPOL2B* (S3C Table).

**S4 Table. Primer sequences used in mutant screening**

**S5 Table. Primer sequences used to amplify *AtPOL2A* and *AtPOL2B* cDNA by RT-qPCR.**

